# CPK3-mediated cytosolic-nuclear translocation of bHLH107 recruits HY5 to regulate the Cu^2+^-triggered upregulation of *ACS8*

**DOI:** 10.1101/2024.09.27.615457

**Authors:** Yue Yu, Haifeng Liu, Xiangsong Chen, Zhaohui Chu

**Affiliations:** State Key Laboratory of Hybrid Rice, Hubei Hongshan Laboratory, College of Life Sciences, Wuhan University, Wuhan 430072, China; State Key Laboratory of Wheat Improvement, College of Agronomy, Shandong Agricultural University, Taian 271018, China; Ezhou Seed Technology Institute of Hubei Province, Ezhou 436043, China

## Abstract

Copper serves as a micronutrient for plant growth and development and has been a key component of copper-based antimicrobial compounds (CBACs) for protection against plant diseases for more than 130 years. We previously revealed that nanomolar-to-micromolar concentrations of Cu^2+^ elicit plant immune responses by activating the expression of the ethylene synthesis rate-limiting enzyme *ACS8*, which is dependent on the promoter copper response element (CuRE) *cis*-element. Here, we genetically confirmed that Cu^2+^-induced resistance to *Pseudomonas syringae* pv. *tomato* (*Pst*) DC3000 is dependent on the CuRE in *ACS8*. Upon screening for CuRE-binding transcription factors, bHLH107, which is required for Cu^2+^-triggered activation of *ACS8* expression and resistance to *Pst* DC3000, was identified via DNA-pull-down and mass spectrometry (MS) assays. Calcium-dependent protein kinase 3 (CPK3) interacts with and phosphorylates bHLH107 at Ser62 and Ser72 to mediate bHLH07 translocation from the cytoplasm into the nucleus, where it interacts with *Arabidopsis* ELONGATED HYPOCOTYL5 (HY5). HY5 directly binds to the G-box and acts as a coactivator to promote bHLH107 binding to the CuRE *cis*-element and to increase transcription of *ACS8* upon Cu^2+^ treatment. Overall, we revealed a CPK3-bHLH107-HY5 module that regulates the Cu^2+^-responsive regulatory network upstream of *ACS8* that is involved in the cytosolic-nuclear translocation of bHLH107.

## INTRODUCTION

Copper is a trace element necessary for plant growth and development. Plants usually exhibit retarded growth and yield loss under conditions of copper deficiency (Burkhead et al., 2009). However, excess copper has toxic effects on plant seed germination, growth and productivity (Mir et al., 2021). Copper is also an effective component of classic plant bactericides and fungicides, which have been used commercially for over 130 years (Lamichhane et al., 2018; Yu et al., 2023a). Copper-based antimicrobial compounds (CBACs), represented by Bordeaux mixture, are widely used to prevent and control plant diseases. Currently, Bordeaux mixture is a commonly used pesticide for fruits and vegetables. For example, copper nanoparticles can effectively protect against *Xanthomonas campestris* pv. *vesicatoria* in tomato (Varympopi et al., 2022). Previously, the mechanism by which CBACs control diseases has relied on two aspects: directly inhibiting the growth of microbial pathogens and the ability of copper hydroxide suspension powder to physically prevent bacterial invasion by covering the surface of plants. Recently, low concentrations of Cu^2+^ have been revealed to trigger plant defense responses and help resist *Pseudomonas syringae* pv. *tomato* (*Pst*) DC3000 infection in *Arabidopsis* (Liu et al., 2015). Moreover, the immune response activated by Cu^2+^ was shown to depend specifically on the ethylene (ET) synthesis rate-limiting enzyme-encoding gene *ACS8* (ACC synthesis 8), whose transcription level rapidly increased after treatment. Furthermore, it activates downstream resistance reactions, including callose deposition, inhibition of the abscisic acid signaling pathway, and activation of the salicylic acid signaling pathway (Zhang et al., 2018; Liu et al., 2020; Yu et al., 2021). An increasing number of studies have provided evidence of Cu^2+^-induced plant immunity. Copper regulates signaling via the SPL9-miR528-AO module to protect against viruses in rice (Yao et al., 2022); additionally, it activates the expression of defense-related (DR) genes to increase resistance to tobacco virus and fungi (Chen et al., 2022a; Guo et al., 2024b). While the copper response element (CuRE) element was shown to mediate signaling responses at the *ACS8* promoter to copper (Zhang et al., 2018), the mechanism of Cu^2+^-mediated *ACS8* expression induction remains unclear.

Transcription factors (TFs) are the master regulators of specific gene expression in plants. Basic helix-loop-helix (bHLH)-containing proteins constitute one of the largest TF families in plants. The typical bHLH domain comprises approximately 60 amino acids, including the basic region at the N-terminus, which is responsible for DNA binding, and the C-terminal HLH region, which promotes protein-protein interactions (Carretero-Paulet et al., 2010). bHLH proteins serve as transcriptional activators and/or suppressors and widely regulate plant growth and development, secondary metabolism, and biotic and abiotic stresses (Gao & Dubos, 2023). For example, bHLH TFs of phytochrome-interacting factors (PIFs) are hubs modulating plant development and growth that can regulate the expression of many downstream genes, integrate multiple signaling pathways, and act as intracellular signal transduction centers (Sun et al., 2024; Leivar & Quail, 2011). As key regulatory factors in secondary metabolism, many bHLH proteins are involved in regulating the synthesis and metabolism of alkaloids and anthocyanins (Gou et al., 2024; Jiang et al., 2024; Guo et al., 2024a). bHLH proteins also confer tolerance or sensitivity to abiotic stresses such as drought, flooding, heat, cold, and salt (Ma et al., 2024; Tian et al., 2024; Wang et al., 2024; Hu et al., 2024; Wang et al., 2022; Zhao et al., 2024; Yan et al., 2023). In addition, they are involved in the plant response to biotic stresses. OsbHLH057 targets AATCA *cis*-elements of the short peptide-encoding gene *Os2H16* to promote rice resistance to *Xanthomonas oryzae* pv. *oryzae* (*Xoo*), *Rhizoctonia solani* and drought (Liu et al., 2022). OsbHLH034 positively regulates rice resistance to *Xoo* by increasing lignin biosynthesis (Onohata & Gomi, 2020). OsbHLH6 promotes rice disease resistance by regulating SA and JA signaling (Meng et al., 2020). The expression of bHLH132 in tomato is highly induced by the XopD effector, which enhances resistance to *X. euvestoria* (Kim & Mudgett, 2019). However, some other bHLH proteins have been reported to negatively regulate plant disease resistance. Nrd1 negatively regulates resistance to *P. syringae* (Zhang et al., 2022b). Notably, the FER-like iron deficiency-induced transcription factor (FIT) undergoes a light-inducible subnuclear partitioning when engaged in protein complexes with itself and with bHLH039 (Trofimov et al., 2024). These bHLH proteins individually or synergistically affect the expression of various target genes, forming complex regulatory networks in plants.

Studies have shown that bHLH proteins ordinarily form homodimers or heterodimers with a wide range of transcriptional regulatory factors, such as bHLH proteins (Toledo-Ortiz et al., 2003), R2R3-MYBs (Yang et al., 2012), bZIPs (Kuras et al., 1997), and epigenetic regulators (Thorstensen et al., 2008). bZIP Met4 and Met28 have been demonstrated to bind with DNA only in the presence of the bHLH protein Cbf1 in yeast. In addition, Met28 could enhance the DNA-binding activity of Cbf1 (Kuras et al., 1997). Additionally, bHLH proteins play important roles in the formation of MYB-bHLH-WD (MBW) complexes associated with anthocyanin synthesis (Yang et al., 2012). The dimerization of bHLH proteins expands their regulatory roles (Carretero-Paulet et al., 2010; Massari and Murre, 2000; Toledo-Ortiz et al., 2003).

ELONGATED HYPOCOTYL 5 (HY5), a bZIP transcription factor, is a master regulator of light-mediated responses. HY5 binds to promoters of approximately 3000 genes containing G-box elements, thereby regulating plant growth and development (Mankotia et al., 2024; Chu et al., 2022; Zhang et al., 2022a; Chen et al., 2022b), photomorphogenesis (Zhang et al., 2024b; Yao et al., 2024; Xiong et al., 2023), transpiration (Kelly et al., 2023), response to biotic and abiotic stress (Yang et al., 2023a; Chen et al., 2021), nutritional homeostasis (Zhang et al., 2014; Mankotia et al., 2023) and more. Some studies have revealed the role of HY5 in the regulation of Cu/Fe homeostasis in response to changing light conditions and Cu/Fe conditions in *Arabidopsis* (Zhang et al., 2014; Mankotia et al., 2023). Moreover, it can positively regulate *Arabidopsis* defense against *Hyaloperonospora arabidopsidis* (Chen et al., 2021). Additionally, HY5 was reported to affect the relative enrichment of H3K9/K14ac on target genes (Guo et al., 2008; Charron et al., 2009; Velanis et al., 2016). Remarkably, a significant portion of light-regulated gene expression depends on GCN5 and TBP-associated factor 1 (TAF1) histone acetyltransferase, which require HY5 (Benhamed et al., 2006; Benhamed et al., 2008; Bertrand et al., 2005).

TFs typically contain nuclear localization signals that are required for their transcriptional regulation functions in the nucleus. However, some TFs are localized in the cytoplasm and are transported into the nucleus only in response to specific signals, including biotic and abiotic stresses. The preformation of TF proteins in the cytoplasm may result in a faster response than *de novo* synthesis (Marathe et al., 2024). Protein modifications such as phosphorylation play important roles in nuclear-cytoplasmic trafficking. BRASSINAZOLE RESISTANT1 (BZR1) interacts with a 14-3-3 protein and is located in the cytoplasm. Receptor for activated C kinase 1 (RACK1) competes interact with BZR1 and promotes trafficking from the cytoplasm to the nucleus after the perception of brassinosteroid (BR) signals (Li et al., 2023). In addition, auxin also induces BZR1 translocation into the nucleus and promotes hypocotyl elongation, which is mediated by mitogen-activated protein kinases (MPKs) and inhibited by 14-3-3 protein (Yu et al., 2023b). Some cytosolic-nuclear trafficking proteins rely on phosphorylation by calcium-dependent protein kinases (CPKs), which constitute the largest subfamily of Ca^2+^ sensors in plants and play important roles in plant growth, development, and response to stress (Zhao et al., 2023; Fan et al., 2023a; Fan et al., 2023b; Bender et al., 2018). CPKs have two domains, N-terminal protein kinase domains and C-terminal CaM-like domains (CLDs), which can activate their kinase domains by directly binding to Ca^2+^ (Harper et al., 2004) and thus transduce signals to induce a series of physiological processes, including abiotic stresses such as drought and high salt levels (Liu et al., 2024; Fan et al., 2023b; Cheng et al., 2022; Rezayian & Zarinkamar, 2023), pollen development (Ranjan et al., 2022), stomatal movement (Schulze et al., 2021; Prodhan et al., 2018; Wang et al., 2019), cell death (Pan et al., 2019; Cui et al., 2020) and the immune response (Zhang et al., 2024a). CPK3 is involved in abscisic acid (ABA) signaling and salicylic acid (SA) signaling, which are associated with stomatal closure in guard cells and herbivory-induced signaling (Mori et al., 2006; Kanchiswamy et al., 2010; Prodhan et al., 2018). In *in vitro* assays, CPK3 has been shown to phosphorylate several TFs, such as ERF1, HsfB2a and CZF1/ZFAR1, which are associated with the transcriptional activation of *DR* genes (Kanchiswamy et al., 2010). Additionally, CPK3 interacts with a 14-3-3 protein to form a complex and phosphorylates 14-3-3 to activate sphingolipid-induced cell death after phytosphingosine treatment in *Arabidopsis* (Lachaud et al., 2013). Peptide flg22 and the effector AvrPphB induce pathogen-associated molecular pattern (PAMP)-triggered immunity (PTI) and effector-triggered immunity (ETI), respectively, to promote resistance to *Pst* DC3000, in which CPK3 plays a critical role by phosphorylating actin-depolymerization factor 4 (ADF4) to govern actin cytoskeletal organization (Lu et al., 2020). Under hypoxic stress, CPK12 is rapidly activated, translocating from the cytoplasm to the nucleus, potentiating plant hypoxia sensing by phosphorylating ERF-VII transcription factors (Fan et al., 2023a). CPK16 interacts with and phosphorylates NADPH oxidase respiratory burst oxidase homolog D (RBOHD) to regulate reactive oxygen species (ROS) production (Yu et al. 2024)

Here, we aimed to identify the Cu^2+^-responsive TFs bound to the CuRE of *ACS8*. We found that a CuRE-binding TF, bHLH107, upregulated *ACS8* transcription and positively regulated the Cu^2+^-induced defense response in *Arabidopsis*. Furthermore, we found that the ability of Cu^2+^ to promote bHLH107 accumulation in the nucleus is dependent on CPK3-mediated phosphorylation of Ser62 and Ser72. In the nucleus, HY5 directly binds to the G-box and acts as a coactivator to promote bHLH107 binding to the CuRE *cis-*element of *ACS8* promoter to trigger downstream immune responses.

## RESULTS

### bHLH107 directly binds to the CuRE *cis-element* in the *ACS8* promoter

Previously, the crucial role of the CuRE *cis*-element in response to Cu^2+^-triggered transcriptional activation was identified in the *ACS8* promoter (Zhang et al., 2018). Here, we elucidated the critical roles of the CuRE in response to Cu^2+^-triggered immunity by introducing *Pro_ACS8_*:g*ACS8* and the CuRE deletion construct *Pro_ACS8ΔCuRE_:gACS8* into *acs8*. First, we examined the bacterial load of *Pst* DC3000 in these lines. We found that the bacterial burden in *acs8* and *Pro_ACS8ΔCuRE_*:*gACS8* (ΔCuRE-2, ΔCuRE-3) plants was significantly greater than that in Col-0 and *Pro_ACS8_*:*gACS8* (WT-2, WT-3) plants under Cu^2+^ treatment (Fig. 1A), indicating that the CuRE is essential for Cu^2+^-induced resistance against *Pst* DC3000. Second, we performed RT-qPCR to test the relative expression levels of *ACS8* in wild-type Col-0, *acs8*, *Pro_AtACS8_*:*gAtACS8* (WT-2, WT-3) and *Pro_ACS8ΔCuRE_*:*gACS8* (ΔCuRE-2, ΔCuRE-3) plants. The results revealed that *ACS8* expression was significantly induced by Cu^2+^ in the Col-0, WT-2, and WT-3 transgenic lines but severely impaired in the ΔCuRE-2 and ΔCuRE-3 lines (Fig. 1B).

**Figure 1.**
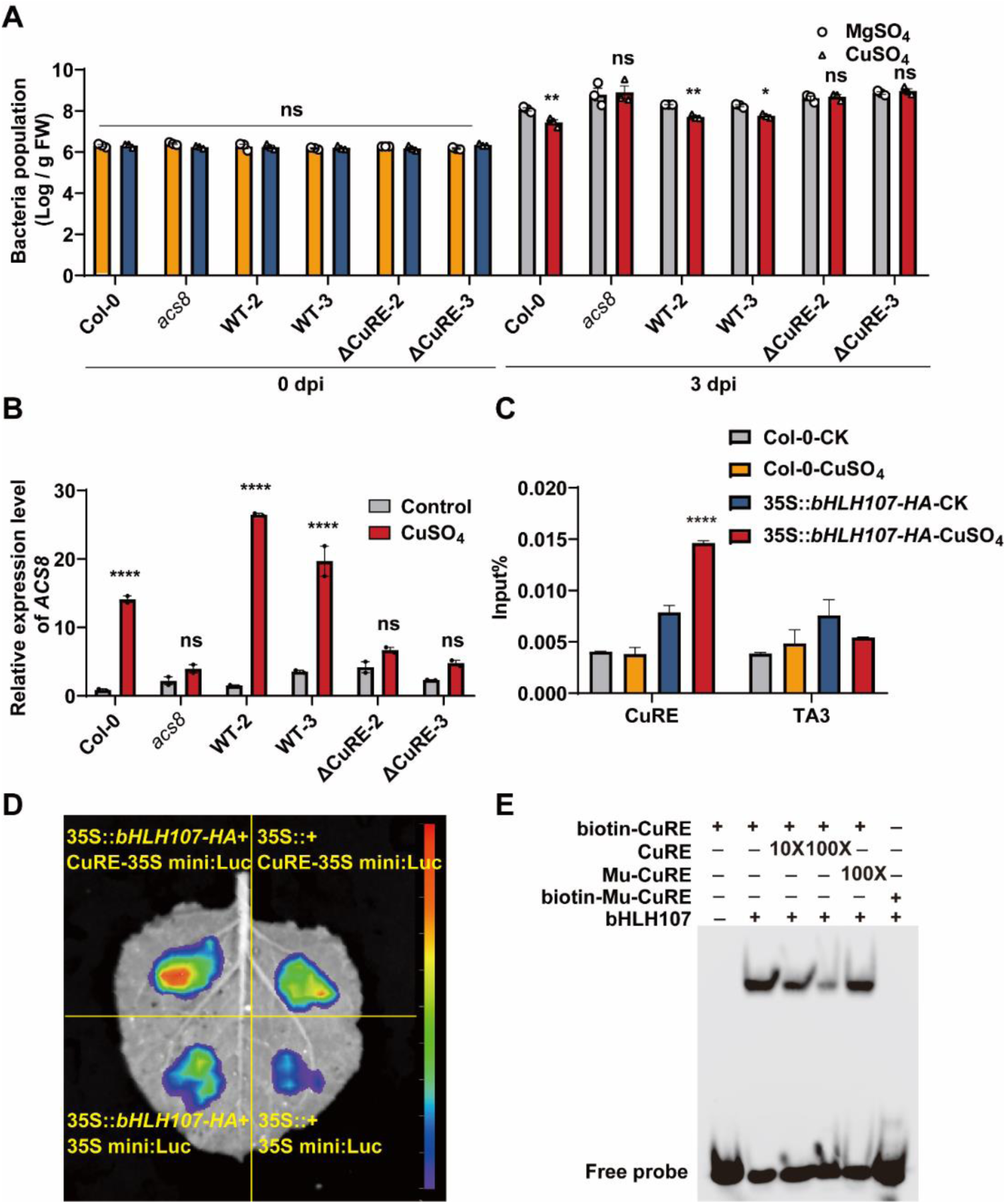
bHLH107 binds to the CuRE *cis-element* in the *ACS8* promoter. (A-B) Bacterial load of *Pst* DC3000 (A) and relative *ACS8* expression (B) in wild-type Col-0, *acs8* mutant, two independent homozygous complemented lines (WT-2, WT-3) and two independent homozygous CuRE mutant complemented lines (ΔCuRE-2, ΔCuRE-3). Four-week-old seedlings were sprayed with MgSO_4_ (100 μM) or CuSO_4_ (100 μM) 4 h before *Pst* DC3000 inoculation. The bacterial population was quantified at 0 dpi and 3 dpi. The error bars represent the means ± SEMs (n = 3). Asterisks indicate significant differences compared with the MgSO_4_-treated plants (two-way ANOVA). The samples were harvested at 2 h after treated with or without CuSO_4_ (100 μM) for RT‒ qPCR assays. The expression levels were normalized to those of *Actin2* (At3g18780). The error bars represent the means ± SEMs (n = 2). Asterisks indicate significant differences compared with the control (two-way ANOVA). (C-E) Identification of bHLH107 bound to the CuRE by ChIP‒qPCR (C), dual-luciferase reporter assays (D) and EMSA (E). Four-week-old Col-0 and 35S::*bHLH107*-*HA* plants were harvested and subjected to ChIP analysis via the anti-HA antibodies shown in Fig. 1A, and the precipitated DNA was recovered and analyzed via qPCR (means ± SEMs; n= 3; two-way ANOVA). Transient coexpression of CuRE-35S mini:Luc or 35S mini:Luc with 35S::*bHLH107*-*HA* or an empty vector in 4-week-old *Nicotiana benthamiana*. Images were taken at 3 d after infiltration, as shown in Fig. 1B. The CuRE sequence (5’-CCAAAAGAAGAAGAAAAACCAAAAGAAGAAGAAAAA-3’) and the Mu-CuRE sequence (5’-CCAAcccccaAAccccAcCCAAcccccaAAccccAc-3’) were designed and labeled with biotin for EMSA, as shown in Fig. 1C.

To identify the transcriptional regulator, DNA pull-down and mass spectrometry (MS) assays were performed to capture the putative TFs bound to CuREs (Table S1). On the basis of the properties of nuclear localization and DNA-binding activity of TFs, eight putative DNA-binding proteins were selected as candidates after filtration of the MS peptides (Fig. S1). At3g56770, which includes a bHLH domain (bHLH107), was shown to activate a dual-luciferase reporter derived from the promoter of *ACS8* after transient co-expression (Figs. 1D, S1). An electrophoretic mobility shift assay (EMSA) was subsequently implemented to demonstrate the CuRE-binding activity of MBP-tagged bHLH107 proteins *in vitro* (Fig. 1E). To examine the bHLH107-CuRE interaction *in vivo*, we constructed a line with 35S-driven constitutive expression of HA-tagged bHLH107 in Col-0 (35S::*bHLH107*-*HA*) plants and performed a chromatin immunoprecipitation-quantitative polymerase chain reaction (ChIP-qPCR) assay. Interestingly, the results revealed that the CuRE was enriched with bHLH107-HA proteins only under Cu^2+^ treatment (Fig. 1C). Together, these results suggest that bHLH107 binds to the key CuRE sequence to activate *ACS8* expression.

### bHLH107 positively regulates Cu^2+^-induced defense responses in *Arabidopsis*

To investigate whether *bHLH107* is required for Cu^2+^-triggered defense responses, we first identified two T-DNA-inserted mutants, GABI_291G03 (*bhlh107-1*/Col-0) and GABI_267B09 (*bhlh107-2*/Col-0), from TAIR-ABRC (Fig. S2) and generated gene editing*-*mutated plants (*bhlh107*) via CRISPR in Col-0 with an early stop codon at the first exon (Fig. S2). We performed RT-qPCR to assess the *ACS8* transcription level and found that the Cu^2+^-induced expression of *ACS8* was significantly reduced in *bhlh107-1, bhlh107-2* and *bhlh107* plants (Fig. 2B, 2D). Moreover, Cu^2+^-induced resistance to *Pst* DC3000 was suppressed in *bhlh107-1*, *bhlh107-2* and *bhlh107* plants compared with that in wild-type Col-0 plants (Fig. 2A, 2C, 2E). Furthermore, Cu^2+^-induced *ACS8* expression and resistance against *Pst* DC3000 were restored in two complemented lines, Com-1 and Com-2 (Fig. 2A, 2C, 2E), which were generated by introducing the *Pro_bHLH107_:gbHLH107-Flag* construct into *bhlh107-1* plants. Collectively, our data demonstrated that bHLH107 positively regulates the Cu^2+^-triggered immune response in *Arabidopsis*.

**Figure 2.**
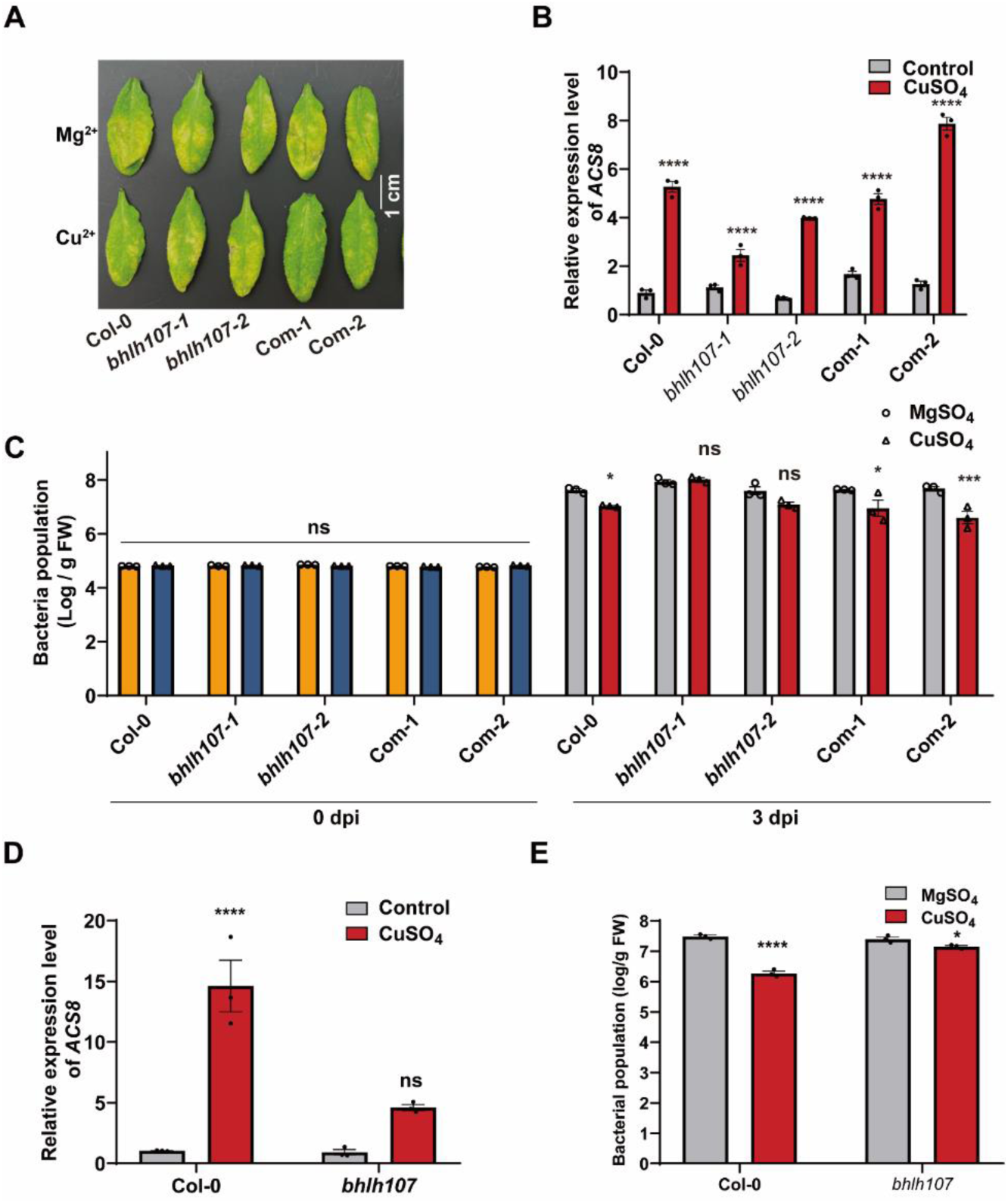
bHLH107 positively regulates Cu^2+^-induced defense responses. (A-C) Disease symptoms (A), relative *ACS8* expression (B) and the bacterial load of *Pst* DC3000 (C) in wild-type Col-0, *bhlh107-1*, *bhlh107-2*, and two independent homozygous complemented lines (Com-1 and Com-2). (D-E) The relative *ACS8* expression (D) and the bacterial load of *Pst* DC3000 (E) in *bhlh107* plants. The samples were harvested at 2 h after treated with or without CuSO_4_ (100 μM) for RT‒qPCR assays. The expression levels were normalized to those of *Actin2*. The error bars represent the means ± SEMs (n = 3). Asterisks indicate significant differences compared with the control (two-way ANOVA). Four-week-old seedlings were sprayed with MgSO_4_ (100 μM) or CuSO_4_ (100 μM) 4 h before *Pst* DC3000 inoculation. The bacterial population was quantified at 0 dpi and 3 dpi. The error bars represent the means ± SEMs (n = 3). Asterisks indicate significant differences compared with the MgSO_4_-treated plants (two-way ANOVA).

### Cu^2+^ stimulates the cytosolic-nuclear translocation of bHLH107

Previous RNA-seq data revealed that bHLH107 expression is upregulated approximately 2.5-fold by Cu^2+^ (Zhang et al., 2018), indicating that bHLH107 may regulate *ACS8* transcription by increasing RNA levels. To determine whether the Cu^2+^-induced defense response in *Arabidopsis* depends on the upregulation of bHLH107 transcription, we examined the bacterial load of *Pst* DC3000 in Col-0 and three independent homozygous overexpression (OE) lines, OE-1, OE-2 and OE-3, constructed by introducing 35S::*bHLH107*-*HA* into Col-0 (Fig S3). We found that all three OE lines showed no significant difference in terms of bacterial load without Cu^2+^ treatment but exhibited slightly significantly inhibited *Pst* DC3000 growth after Cu^2+^ treatment (Fig. 3A). Moreover, the *ACS8* transcript level in the three OE lines was not significantly greater than that in Col-0 plants in the control conditions, but there was a small significant increase in *ACS8* expression after Cu^2+^ treatment (Fig. 3B). Collectively, these data implied that bHLH107 activation of Cu^2+^-induced resistance may not be dependent on increased transcription and that protein modification may play a major role in regulating the Cu^2+^-induced defense response in *Arabidopsis*.

**Figure 3.**
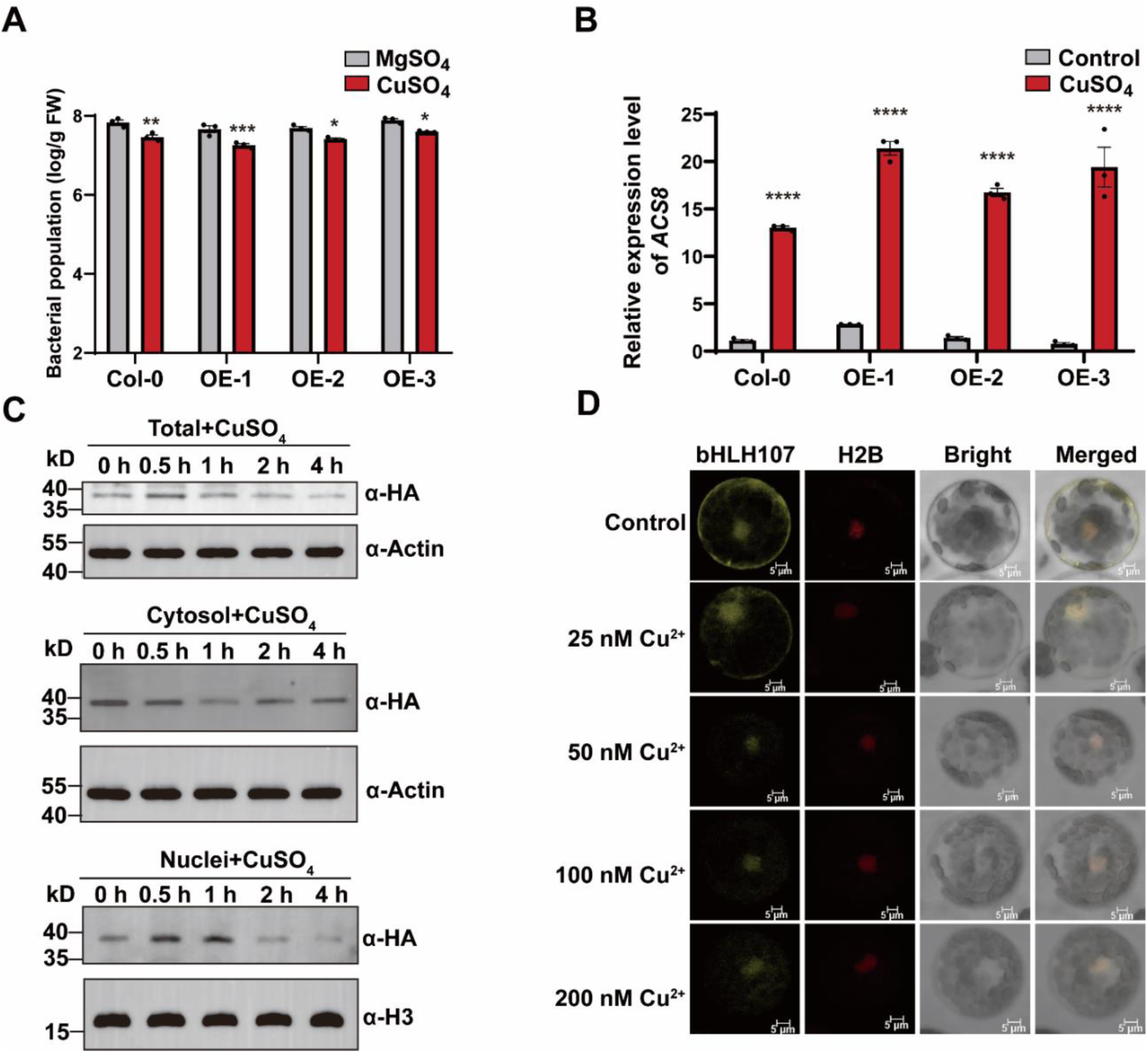
Cu^2+^ activates the cytosolic-nuclear translocation of bHLH107. (A-B) Bacterial load of *Pst* DC3000 (A) and the relative *ACS8* expression (B) were measured for 4-week-old Col-0 and OE-1, OE-2, and OE-3 35S::*bHLH107*-*HA* plants. The methods used were previously described in Fig. 2 (C) Western blot showing the abundance of bHLH107 protein in mixed samples of OE-1, OE-2, and OE-3 plants treated with 100 μM CuSO_4_. The total, cytosolic and nuclear proteins were separated from the leaf samples. Anti-actin and anti-H3 antibodies were used for standard markers for total, cytosolic and nuclear proteins, respectively. (D) Subcellular localization of bHLH107-YFP in response to different concentrations of CuSO_4_. H2B-mCherry was used as a nuclear localization marker. Scale bars, 5 μm.

To further investigate the subcellular localization of bHLH107, we used a cell fractionation assay to investigate whether CuSO_4_ could redistribute the abundance of bHLH107 at different subcellular locations. Interestingly, the abundance of bHLH107 protein was lower than that of total proteins and cytoplasmic proteins at 2 h and 1 h, respectively, while bHLH107 protein accumulated in the nucleus at 0.5 h and 1 h after treatment with CuSO_4_ (Fig. 3C). To further investigate this result, we generated a *ProbHLH107:gbHLH107*-YFP construct and introduced it into Col-0 protoplasts and detected bHLH107-YFP signals via confocal microscopy. We observed that bHLH107-YFP was located mainly in the cytoplasm and nucleus in the control condition, while it was located predominantly in the nucleus after treatment with CuSO_4_ (Fig. 3D). These results suggest that Cu^2+^ treatment could stimulate bHLH107 translocation from the cytoplasm into the nucleus. Moreover, we detected increased phosphorylation of bHLH107 after Cu^2+^ treatment by using a Phos-tag SDS-PAGE assay (Fig. S4). Together, our data indicate that Cu^2+^ stimulates the cytosolic-nuclear translocation of bHLH107, which is associated with its phosphorylation.

### CPK3 interacts with bHLH107 and participates in Cu^2+^-triggered immunity

To elucidate the mechanism of nuclear enrichment of bHLH107, an IP-MS assay was performed to identify potential bHLH107-binding proteins. We found that calcium-dependent protein kinase 3 (CPK3), a Ca^2+^ sensor, was a strong candidate for interaction with bHLH107 (Table S2). To confirm the interaction between CPK3 and bHLH107, luciferase complementation imaging (LCI), pull-down and semi-*in vivo* coimmunoprecipitation (co-IP) assays were performed (Fig. 4). Co-expressing CPK3-nLuc and cLuc-bHLH107 activated complete luciferase activity through physical interaction, but co-expressing CPK3-nLuc with cLuc-Actin or cLuc-bHLH107 with nLuc-Actin did not (Fig. 4A). Additionally, the MBP-tagged bHLH107 proteins were immunoprecipitated by GST-tagged CPK3 proteins but not the control GST tag (Fig. 4B). Moreover, a semi-*in vivo* co-IP assay revealed that the MYC-tagged CPK3 proteins expressed in *N. benthamiana* were immunoprecipitated via prokaryotically expressed MBP-tagged bHLH107 proteins but not the MBP tag (Fig. 4C). Overall, we concluded that bHLH107 physically interacts with CPK3.

**Figure 4.**
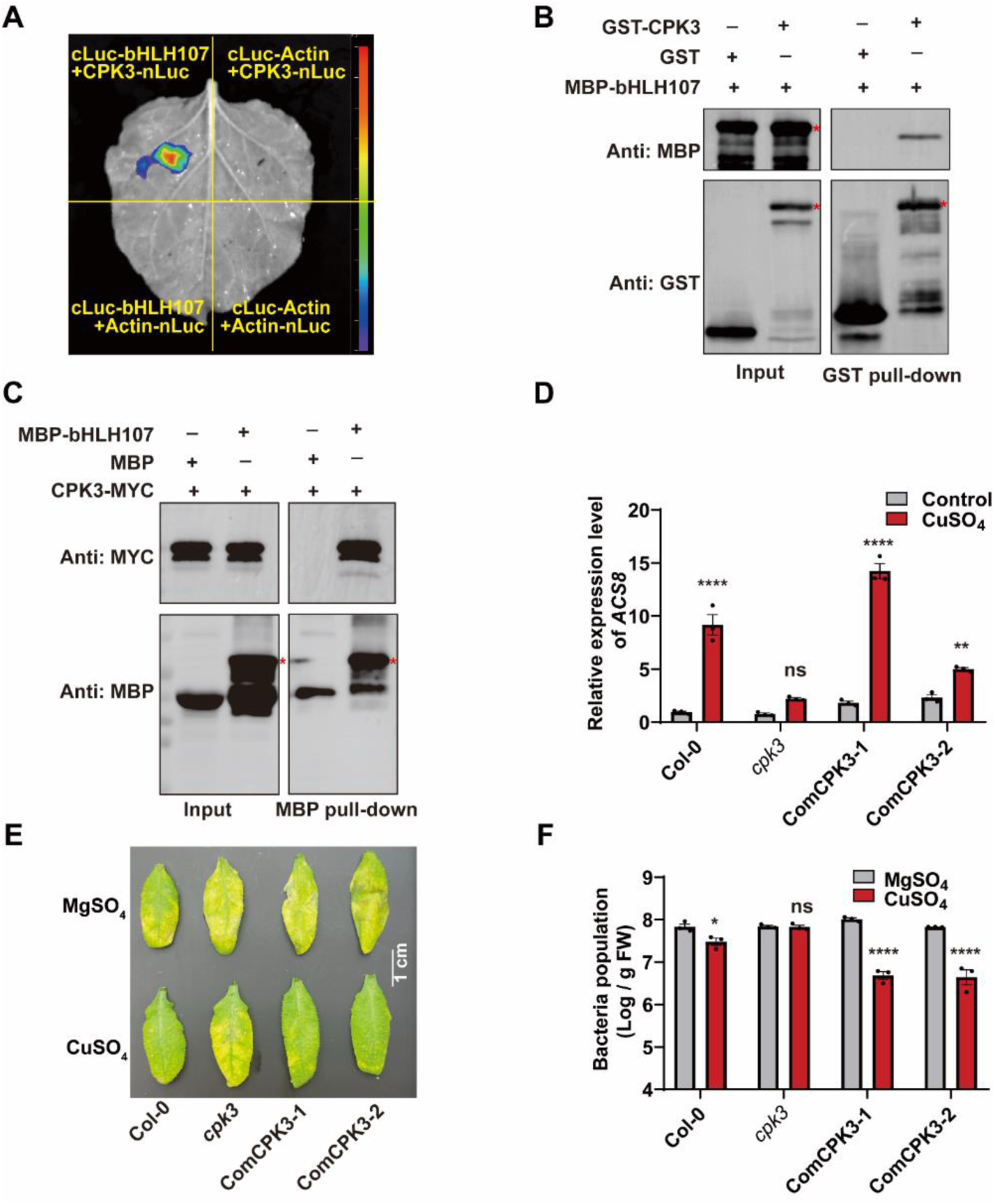
CPK3 interacts with bHLH107 and is required for Cu^2+^-induced resistance. (A-C) The interaction of CPK3 and bHLH107 was assessed with LCI (A), pull-down (B) and semi-*in vivo* co-IP (C) assays. Transiently coexpression of cLuc-bHLH107 with CPK3-nLuc was used as the experimental group, and cLuc-bHLH107 or CPK3-nLuc was coexpressed with Actin-nLuc or cLuc-Actin as the native control in 4-week-old *N. benthamiana*. Images were obtained at 36 h after infiltration, as shown in Fig. 4A. Prokaryotically expressed GST-tagged CPK3 but not GST alone could pull down MBP-tagged bHLH107 *in vitro*. The precipitated fractions were analyzed with anti-GST and anti-MBP antibodies, as shown in Fig. 4B. semi-*in vivo* co-IP was performed with plants expressing MYC-tagged CPK3 and prokaryotically expressed MBP-tagged bHLH107. (D-F) The relative *ACS8* expression (D), disease symptoms (E) and bacterial load (F) in Col-0, *cpk3*, and two independent homozygous complementation lines (ComCPK3-1 and ComCPK3-2). The methods used were previously described in Fig. 2.

To investigate whether *CPK3* is required for Cu^2+^-induced resistance, we identified a T-DNA-inserted mutant, *cpk3* (Salk_107620), which has a T-DNA insertion into the 5’UTR of CPK3 and exhibits almost no CPK3 protein expression (Mehlmer et al., 2010). First, we found that Cu^2+^-induced expression of *ACS8* was significantly reduced in *cpk3* plants (Fig. 4D), but inducible expression was rescued in two independent homozygous lines, *Pro_CPK3_:gCPK3*-*Flag-1*/*cpk3* (ComCPK3-1) and *Pro_CPK3_:gCPK3*-*Flag-2*/*cpk3* (ComCPK3-2). Moreover, Cu^2+^-induced resistance to *Pst* DC3000 was abrogated in *cpk3* plants but restored in ComCPK3-1 and ComCPK3-2 plants (Fig. 4E). This finding was consistent with the measured *Pst* DC3000 burden in Col-0, *cpk3,* ComCPK3-1 and ComCPK3-2 plants (Fig. 4F). Thus, we concluded that CPK3 interacts with bHLH107, which is required for the Cu^2+^-induced activation of *ACS8* expression and defense against *Pst* DC3000.

### CPK3 phosphorylates bHLH107 at Ser62 and Ser72

To investigate the potential link between bHLH107 and CPK3, we performed an *in vitro* kinase activity assay and a phos-tag gel immunoblotting assay. The results showed that CPK3 could phosphorylate bHLH107 (Fig. 5A). Phosphorylated bHLH107 was analyzed by mass spectrometry to identify the phosphorylated residues. Four serine/threonine residues (Ser72, Thr164, Thr169, and Ser204) of bHLH107 were found to be phosphorylated (Fig. S5, Table S3). The phosphorylation sites were confirmed via a dual-luciferase reporter system in *N. benthamiana* leaves coexpressing CuRE-35S mini::Luc with 35S:*bHLH107*, which was constructed via site-directed mutagenesis of each serine/threonine residue to the nonphosphorylatable residue alanine (S72A, T164A, T169A, and S204A). The results revealed that bHLH107 with the Ser72 residue mutation could not activate the expression of CuRE-35S mini::Luc, which indicated that bHLH107 Ser72 may be important for activating *ACS8* transcription (Figs. S6, 5B).

**Figure 5.**
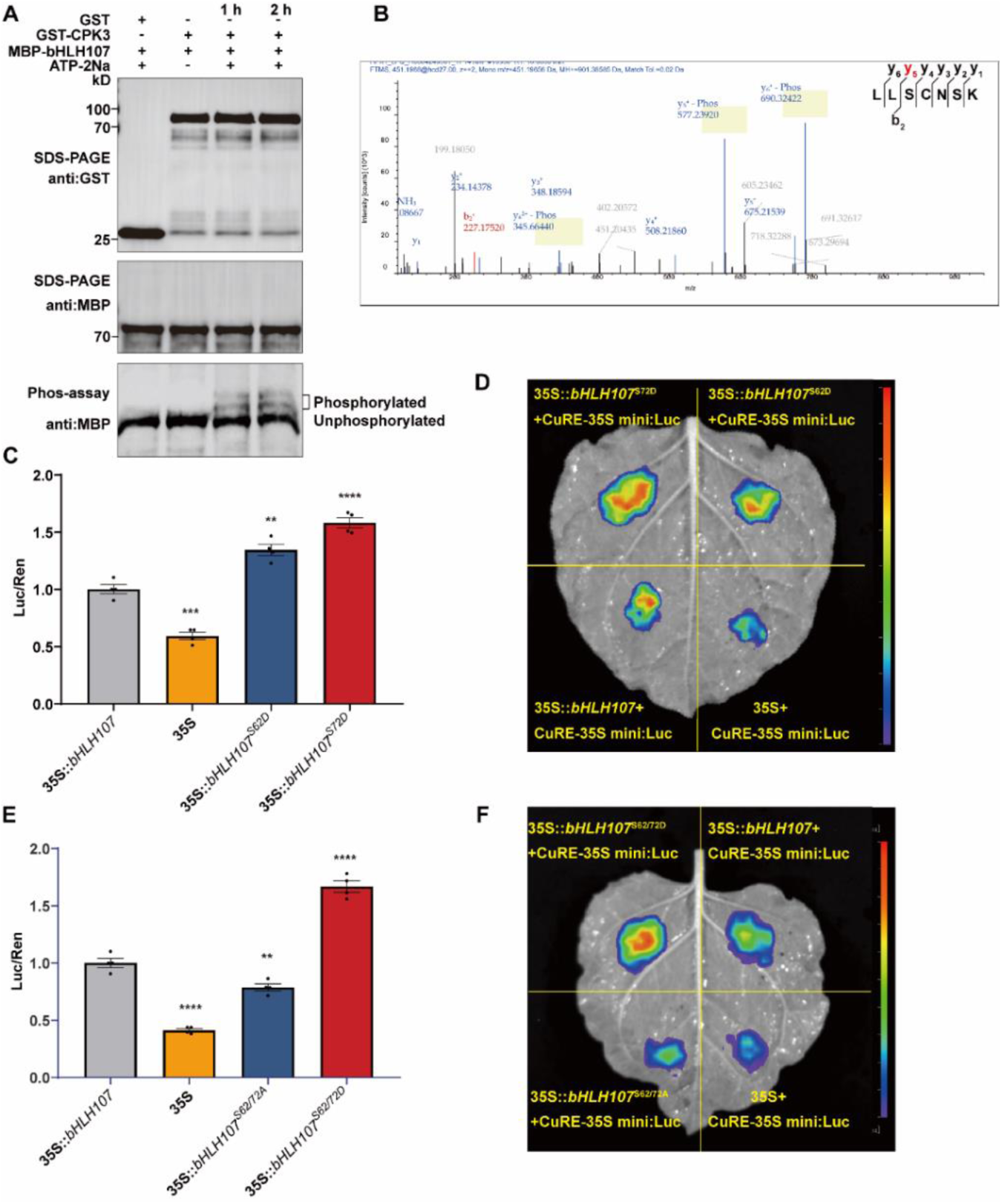
CPK3 phosphorylates serine residues 62 and 72 of bHLH107 to facilitate the transcriptional activation function of bHLH107. (A) *In vitro* kinase assay showing bHLH107 phosphorylation by CPK3. Prokaryotically expressed GST-CPK3 and MBP-bHLH107 were incubated in kinase buffer in the presence or absence of ATP-2Na. Phosphorylated bHLH107 was detected via Phos-tag gel immunoblotting with an anti-MBP antibody. (B) Identification of the phosphorylated residue(s) of bHLH107 via mass spectrometry (C-D) Relative luciferase activity (C) and dual-luciferase reporter assays (D) showing that Ser62 and Ser72 play important roles in activating the expression of the *Reporter* gene. (E-F) Relative luciferase activity (E) and dual-luciferase reporter assays (F) showing that S62/72D and S62/72A play bidirectional roles in activating the expression of the *Reporter* gene. The *Reporter* gene is CuRE-35S mini:Luc, means ± SEMs; n = 3; two-way ANOVA.

Previous studies revealed a specific CPK-binding motif in plant systems, R/KXXS/T (X represents any amino acid) (Furihata et al., 2006). There are two putative CPK-binding motifs, i.e., residues 59-62 (RINS) and 69-72 (KLLS), in bHLH107. The KLLS motif is binds to Ser72, which is phosphorylated by CPK3 (Fig. 5B). Additionally, the other conserved motif, RINS (residues 59-62), was identified on bHLH107, which implied that Ser62 may also be phosphorylated by CPK3. For verification, we introduced phosphomimetic mutations of the target residues (S62D, S72D) to construct a dual-luciferase reporter system in *N. benthamiana* leaves in the presence of co-expressed CuRE-35S mini::Luc. Compared with 35S::*bHLH107*, which increased luciferase expression, the co-expression of the S62D and S72D mutants further increased the expression of the reporter gene (Fig. 5C, D). Alternatively, the dual-phospho-defective mutation protein *bHLH107^S62/72A^* lowered the enzyme activity of CuRE-35S mini::Luc compared with that with 35S::*bHLH107*. However, the dual-phospho-mimetic mutation protein *bHLH107^S62/72D^* increased the enzyme activity of CuRE-35S mini::Luc compared with that with 35S::*bHLH107* (Fig. 5E, F). These results indicated that the putative phosphorylation sites at Ser62 and Ser72 of bHLH107 are targeted sites of CPK3 and are associated with *ACS8* activation.

### Ser62 and Ser72 are required for the nuclear accumulation of bHLH107

We hypothesized that Cu^2+^ induces the cytosolic-nuclear translocation of bHLH107 by phosphorylation; therefore, we tested whether the Ser62 and Ser72 residues determine bHLH107 trafficking. *Pro_bHLH107_:gbHLH107^S62/72A^*-YFP and *Pro_bHLH107_:gbHLH107^S62/72D^*-YFP were transformed into Col-0 protoplasts, and YFP signals were detected via confocal microscopy. bHLH107^S62/72D^ proteins were located mainly in the nucleus, whereas bHLH107^S62/72A^ proteins were located in both the cytoplasm and nucleus in the presence or absence of CuSO_4_. Importantly, Cu^2+^ failed to induce nuclear enrichment of bHLH107^S62/72A^ (Fig. 6A). We also transformed *Pro_bHLH107_:gbHLH107*-YFP, *Pro_bHLH107_:gbHLH107^S627A^*-YFP, and *Pro_bHLH107_:gbHLH107^S627D^*-YFP into *cpk3* mutant protoplasts. Both the bHLH107 and bHLH107^S62/72A^ proteins were located mainly in the cytoplasm and nucleus in the presence or absence of CuSO_4_, and the bHLH107^S62/72D^ proteins were located mainly in the nucleus (Fig. 6B). These results suggest that the Cu^2+^-induced enrichment of bHLH107 in the nucleus depends on phosphorylation of the Ser62 and Ser72 residues by CPK3.

**Figure 6.**
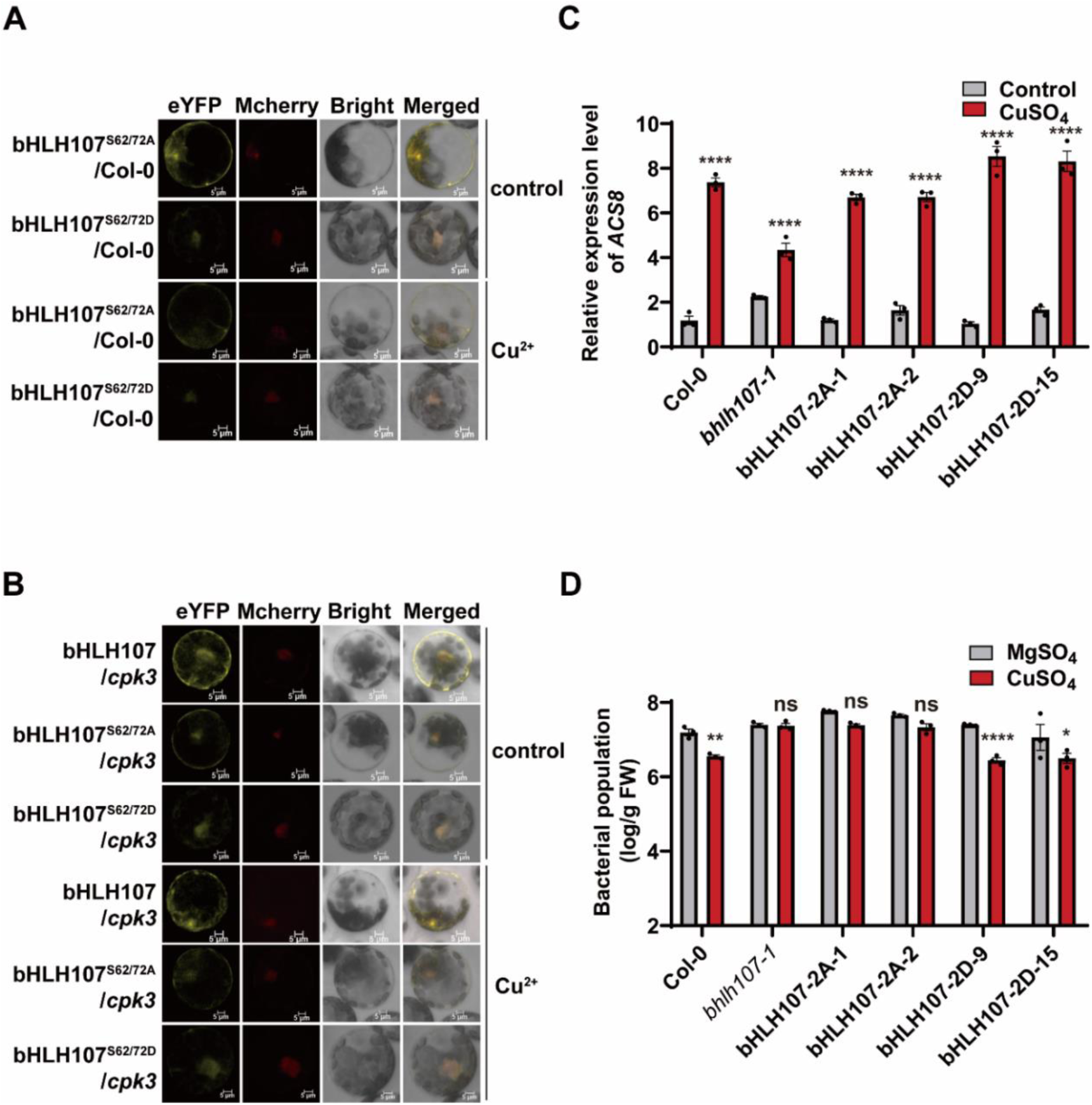
Phosphorylated Ser62 and Ser72 residues are required for the nuclear accumulation of bHLH107 and the Cu^2+^-induced defense response. (A) Subcellular localization of bHLH107^S62/72A^ and bHLH107^S62/72D^ transformed into Col-0 protoplasts in response to 100 nM CuSO_4_. (B) Subcellular localization of bHLH107, bHLH107^S62/72A^ and bHLH107^S62/72D^ transformed into *cpk3* protoplasts in response to 100 nM CuSO_4_. H2B-mCherry was used as a nuclear localization marker. Scale bars, 5 μm. (C-D) The relative transcription of *ACS8* (C) and the bacterial load of *Pst* DC3000 (D) in 4-week-old Col-0, *bhlh107-1*, dual-phospho-defective mutant lines bHLH107-2A (−1, −2) and dual-phospho-mimetic mutant lines bHLH107-2D (−9, −15). The methods used were previously described in Fig. 2.

CPK3-mediated phosphorylation of the Ser62 and S72 residues in bHLH107 indicated that these residues could be crucial for Cu^2+^-induced resistance to *Pst* DC3000. For validation, we first constructed *Pro_bHLH107_:gbHLH107^S62/72A^-Flag* (bHLH107-2A) and *Pro_bHLH107_:gbHLH107^S62/72D^-Flag* (bHLH107-2D) and then transformed them into *bhlh107-1* plants to generate phospho-mutant complemented lines. We performed RT-qPCR to examine *ACS8* transcription levels and found that the Cu^2+^-induced *ACS8* expression was reduced in the dual-phospho-defective mutant line bHLH107-2A (Fig. 6C). Correspondingly, Cu^2+^ could upregulate *ACS8* transcription in the dual-phospho-mimetic mutant line bHLH107-2D (Fig. 6C). Moreover, the bacterial load of *Pst* DC3000 in *bhlh107-1* and bHLH107-2A plants (1, 2) was significantly greater than that in wild-type Col-0 plants, and Cu^2+^-mediated induction of resistance was suppressed. In contrast, bHLH107-2D plants showed decreased susceptibility to *Pst DC3000* (9,15) (Figs. 6D, S7). The above results indicated that the Ser62 and Ser72 residues of *bHLH107* participate in resistance to *Pst* DC3000. In summary, the Ser62 and Ser72 residues are required for the cytosolic-nuclear translocation of bHLH107 and the Cu^2+^-induced defense response.

### HY5 interacts with bHLH107 in nuclei and participates in Cu^2+^-triggered immunity

bHLH107 was previously found among HY5-immunoprecipitated proteins and was shown to interact with HY5 via yeast two-hybrid (Y2H) assay (Gong et al., 2008). Interestingly, HY5 is also involved in copper homeostasis (Zhang et al., 2014). Thus, HY5 may be involved in the transcriptional regulation of Cu^2+^-induced defense responses mediated by bHLH107. First, we validated the interactions between bHLH107 and HY5 via co-IP (Fig. 7A), LCI (Fig. 7B) and pull-down (Fig. 7C) assays. Furthermore, we obtained the *HY5* constitutive expression line 35S::*HY5*-*GFP* in the Col-0 background and the mutant line *ks50* in the WS-0 background (Shi, et al., 2024; Oyama et al., 1997). Interestingly, the Cu^2+^-induced expression level of *ACS8* was not significantly altered in the 35S::*HY5-GFP* and *ks50* lines compared with that in wild-type Col-0 and WS-0 plants, respectively, without Cu^2+^ treatment, whereas the induced expression level of *ACS8* was reduced in the *ks50* line, and there was no obvious effect in the 35S::*HY5-GFP* line at 2 h after treatment with Cu^2+^ (Fig. 7D, 7E). Consistent with the positive regulation of *ACS8* expression, 35S::*HY5-GFP* plants presented increased resistance to *Pst* DC3000 compared with that of Col-0 plants under both MgSO_4_ and CuSO_4_ treatment (Fig. 7F). However, the bacterial load in *ks50* plants was significantly greater than that in WS-0 plants under Cu^2+^ treatment (Fig. 7G), indicating that the loss-of-function *HY5* mutation impaired the ability of Cu^2+^ to induce a defense response against *Pst* DC3000 (Fig. 7F, G). Thus, our results demonstrate that HY5 physically interacts with bHLH107 to regulate Cu^2+^-induced defense responses.

**Figure 7.**
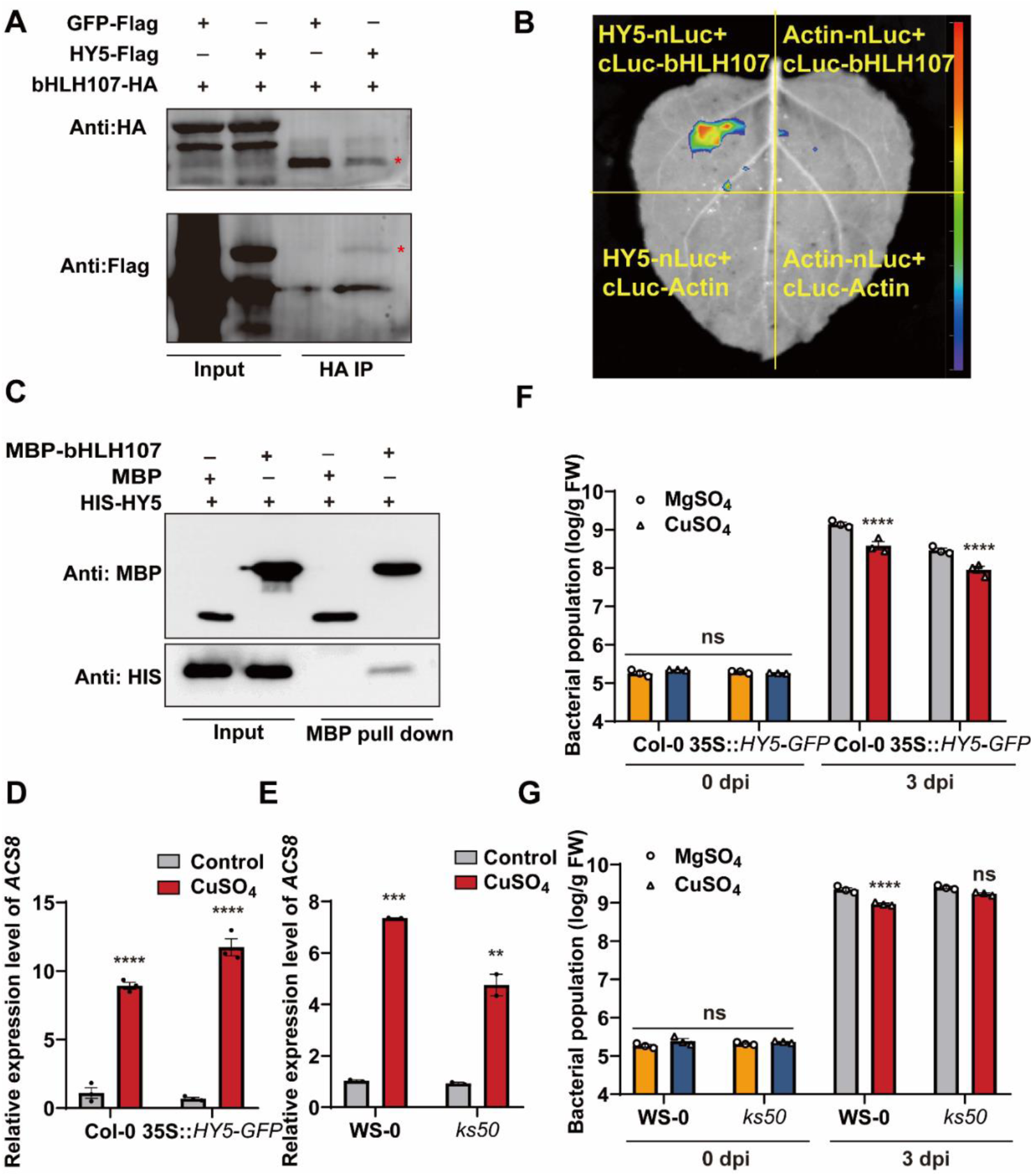
HY5 physically interacts with bHLH107 and participates in Cu^2+^-induced defense responses. (A-C) HY5 interacts with bHLH107, as determined via co-IP (A), LCI (B) and pull-down (C) assays. HA-tagged bHLH107 was coexpressed with Flag-tagged HY5 in 4-week-old *N. benthamiana* as the experimental group and coexpressed with Flag-tagged GFP as a negative control. Total protein was purified at 36 h after infiltration and subsequently incubated and precipitated with Flag-tagged beads. Western blots were performed with anti-Flag and anti-HA antibodies, as shown in Fig. 7A. cLuc-bHLH107 was coexpressed with Actin-nLuc, cLuc-Actin with HY5-nLuc, or cLuc-Actin with Actin-nLuc as the native control in 4-week-old *N. benthamiana*, as shown in Fig. 7B. Images were taken at 36 h after infiltration. His-tagged HY5 was incubated with immobilized MBP or MBP-tagged bHLH107; subsequently, the precipitated fractions were analyzed with anti-His and anti-MBP antibodies, as shown in Fig. 7C. (D-E) RT‒qPCR assays showing the relative expression of *ACS8* in the wild-type Col-0 and *HY5* over-expression transgenic lines *35S::HY5-GFP*/Col-0 (D) and in the wild-type WS-0 and HY5 mutant plants *ks50*/WS-0 (E). Methods were performed according to the protocols described above. (F-G) Bacterial load of *Pst* DC3000 in Col-0 *35S::HY5*-*GFP*/Col-0 plants (F) and in WS-0 and *ks50*/WS-0 plants (G) at 0 dpi and 3 dpi. *Arabidopsis* plants were sprayed with MgSO_4_ (100 μM) or CuSO_4_ (100 μM) 4 h before *Pst* DC3000 infiltration. The error bars represent the means ± SEMs (n = 3). Asterisks indicate significant differences compared with the MgSO_4_-treated plants (two-way ANOVA).

### HY5 serves as a coactivator to upregulate *ACS8* expression

Promoter analyses have revealed that the G-box is bound by HY5 *in vitro* (Chattopadhyay et al., 1998). There are seven putative G-boxes located on the *ACS8* promoter (Fig. 8A). To explore whether HY5 can bind to the *ACS8* promoter, we first performed a yeast one-hybrid (Y1H) assay. The promoter sequence was separated into 8 fragments from “a” to “h” for the Y1H binding activity assay (Fig. 8A, 8B). The results revealed that fragments “b” and “c” in the *ACS8* promoter bound to HY5 (Fig. 8B). Additionally, EMSAs revealed that both G-box1 on the “b” fragment and G-box2 on the “c” fragment bound with HY5 (Fig. 8C, 8D). To explore the role of HY5 in bHLH107 binding to CuREs *in vivo*, we generated 35S::*bHLH107*/*ks50* and 35S::*bHLH107*/WS-0 transgenic lines and performed a ChIP-qPCR assay to test the binding activity. The qPCR results indicated that the Cu^2+^-induced increase in CuRE binding was eliminated in 35S::*bHLH107*/*ks50* compared with 35S::*bHLH107*/WS-0 plants (Fig. 8E). These results suggest that HY5 cooperates with bHLH107 and binds to G-boxes and CuREs to facilitate *ACS8* expression.

**Figure 8.**
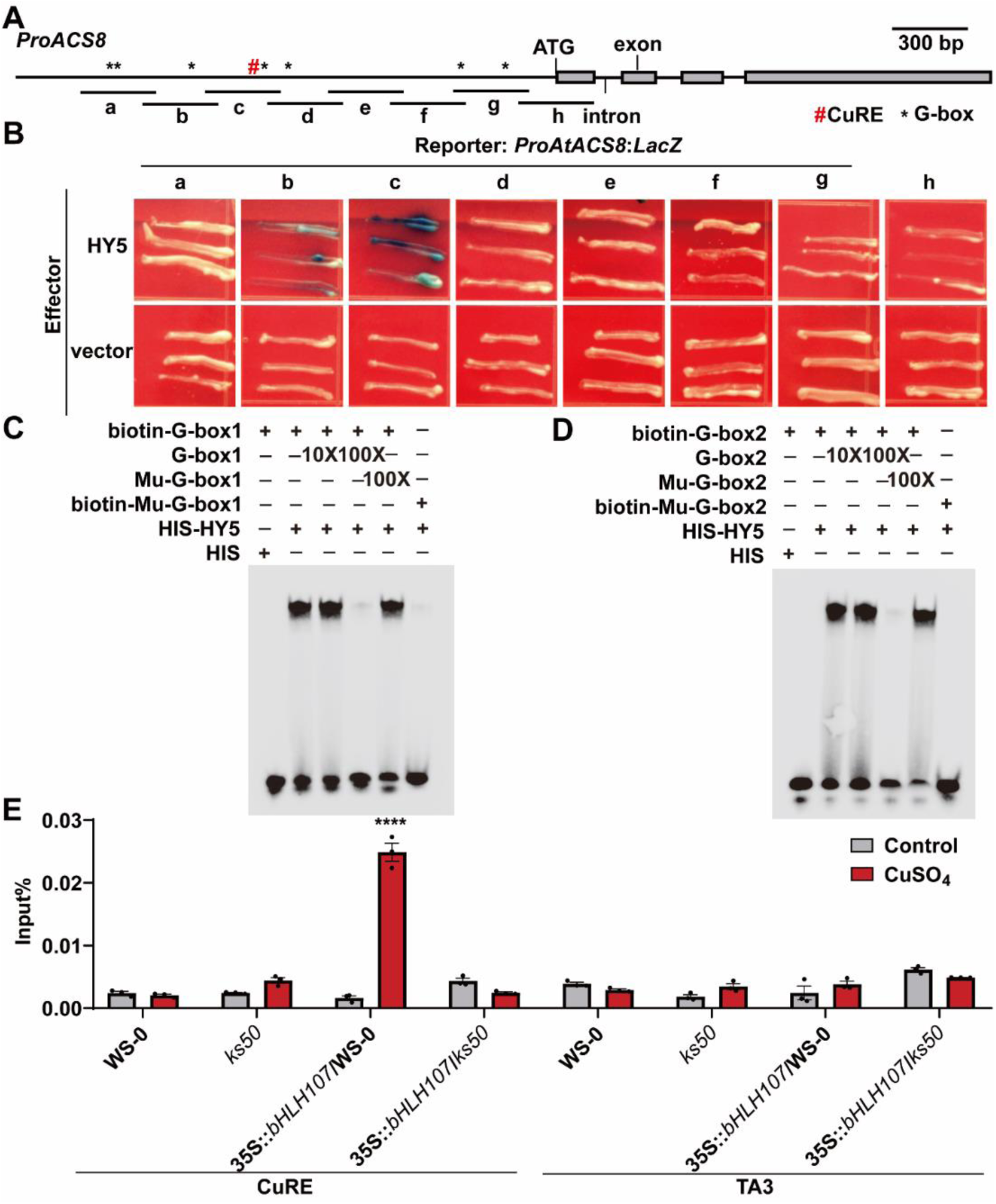
HY5 serves as a coactivator by binding to the G-box adjacent to the CuRE to upregulate *ACS8* expression. (A) Illustration of *ACS8* promoter fragments used for yeast one-hybrid (Y1H) assays. The adenine residue of the translation initiation site (ATG) was designated position +1, and the promoter sequence was segmented into nine DNA fragments named “a” to “h” for Y1H assays. (B) Y1H data showing that HY5 binds to G-box1 and G-box2 of the *ACS8* promoter. HY5 or the vector control and various *LacZ* reporters were cotransformed into the yeast strain EGY48. (C-D) EMSAs. The G-box1 sequence (5’-AAACTGAAAAACACGTTGTTAGTTTATATG-3’), Mu-G-box1 sequence (5’-AAACTGAAAAACtttcTGTTAGTTTATATG-3’), Mu-G-box2 sequence (5’-GGTATGTTCCGACtttcTTCCGGCGAGAAC-3’) and the G-box2 sequence (5’-GGTATGTTCCGACACGTTTCCGGCGAGAAC-3’) were designed and labeled with biotin. (E) ChIP‒qPCR assays showing that bHLH107 binding to the CuRE sequence is dependent on HY5 *in vivo*. Four-week-old 35S::*bHLH107*/WS-0 and 35S::*bHLH107*/*ks50* plants were harvested and subjected to ChIP analysis via anti-HA antibodies, and the precipitated DNA was recovered and analyzed via qPCR assays (means ± SEMs; n= 3; two-way ANOVA).

## DISCUSSION

Cu^2+^, an essential trace element in plants, participates in multiple vital processes, such as photosynthesis and respiration (Drepper et al., 1996; Soriano et al., 1996). For plant disease management, it has also been a key component of CBACs for over thirteen decades (Lamichhane et al., 2018; Yu et al., 2023a). However, overuse of CBACs has resulted in long-term exposure to high concentrations of Cu^2+^, which has led to the emergence of many Cu^2+^-resistant bacterial strains (Behlau et al., 2013; Voloudakis et al., 2005). Researchers have reported that the role of Cu^2+^ in controlling pathogenic microorganisms relies not only on sterilizing activity but also on stimulating plant immune responses (Liu et al., 2015, 2020; Yao et al., 2022; Chen et al., 2022a; Guo et al., 2024b). In *Arabidopsis*, the immune response needed to resist *Pst* DC3000 is activated by Cu^2+^ and is specifically dependent on the CuRE *cis*-element located in the *ACS8* promoter, which rapidly activates ET signaling after treatment (Liu et al., 2015; Zhang et al., 2018). However, the mechanism of Cu^2+^-mediated induction of *ACS8* expression remains unclear. Here, we identified the CuRE-binding TF bHLH107 and systematically elucidated the Cu^2+^-mediated immune response pathway upstream of *ACS8.* The proposed pathway is shown in Fig. 9. *Arabidopsis* perceives Cu^2+^ and activates CPK3, which interacts with and phosphorylates bHLH107 to mediate its translocation into the nucleus; subsequently, bHLH107 cooperates with HY5, increase the transcription level of *ACS8* (Fig. 9).

**Figure 9.**
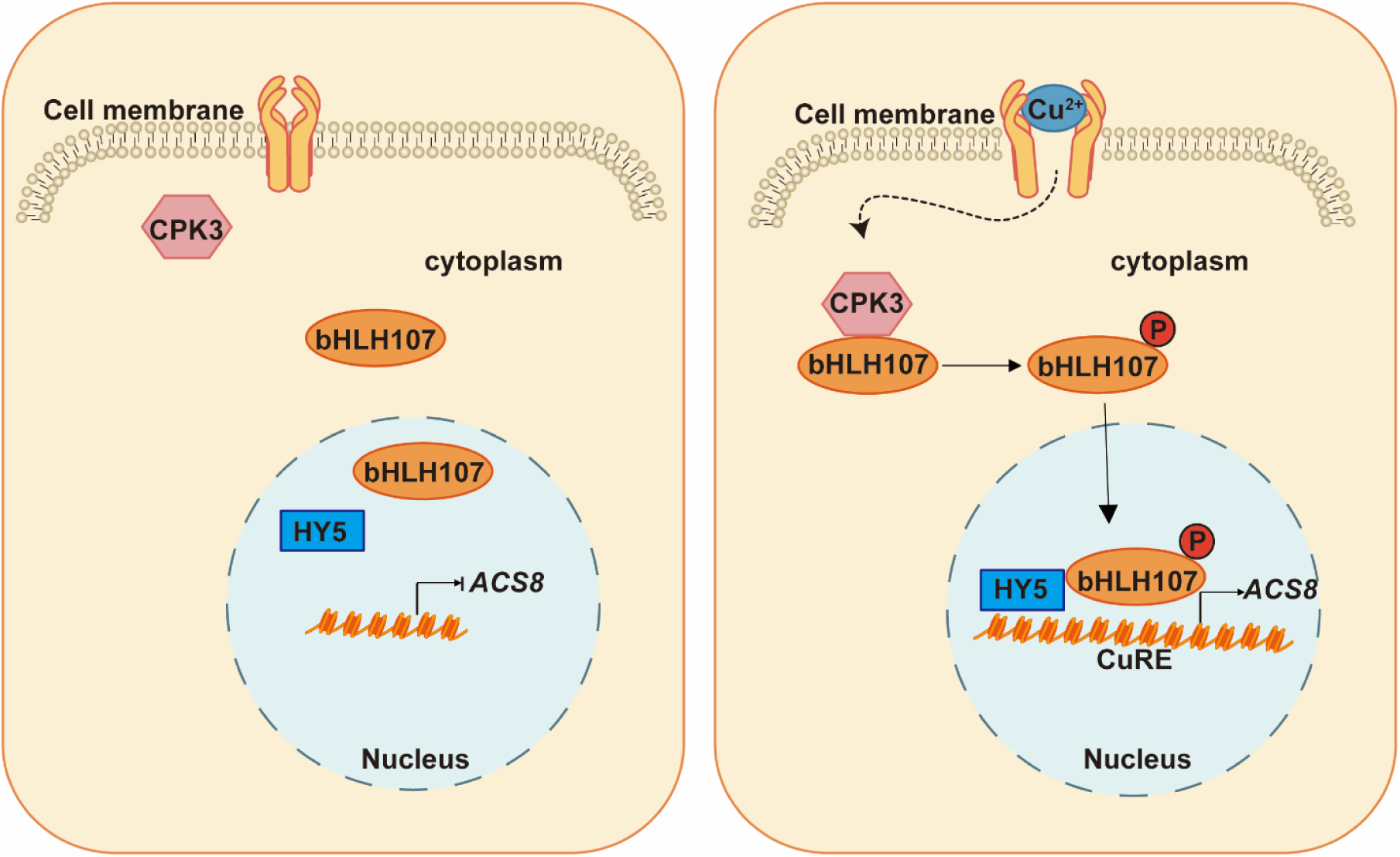
A model for the transcriptional module of copper-activated *ACS8* expression in *Arabidopsis*.

Previous studies have indicated that copper can elicit an immune response in rice and tobacco (Yao et al., 2022; Chen et al., 2022a; Guo et al., 2024b). According to our results together with our previous findings, low concentrations of Cu^2+^ can trigger a plant defense response by regulating the CuRE *cis*-element in the *ACS8* promoter, which is rapidly upregulated at the transcriptional level after treatment (Liu et al., 2015; Zhang et al., 2018). We demonstrated that bHLH107 was translocated from the cytoplasm into the nucleus and cooperated with HY5 to bind the CuRE to upregulate *ACS8* transcription. This finding provides insight into the Cu^2+^-triggered immunity upstream of *ACS8*. Second, as a TF, bHLH107 was only found to interact with HY5 and be activated by chitin (Libault et al., 2007; Gong et al., 2008); however, the regulatory network involved is unknown. In this study, we found that Cu^2+^ could induce *ACS8* transcription in both Col-0 and two independent bHLH107 complemented lines, Com-1 and Com-2. Thus, we revealed a novel function of bHLH107 in the Cu^2+^-induced defense response in *Arabidopsis*. However, the induction of *ACS8* transcription by Cu^2+^ was partially alleviated in both the T-DNA-inserted *bhlh107-1* mutant and the *bhlh107* knockout line (Fig. 2C, S2), suggesting that bHLH107 is not a unique TF regulating Cu^2+^-induced *ACS8* transcription. Other TFs may bind to the CuRE and regulate *ACS8* transcription. Consistent with these findings, by using a dual-luciferase reporter system in *N. benthamiana* leaves, we confirmed that At1g61730, At4g16430 and At3g22900 could also activate reporter gene expression after transient coexpression with CuRE-35S mini:Luc (Fig. S1).

Some TFs are localized in the cytoplasm and transported into the nucleus only in response to specific signals, including biotic and abiotic stresses. This phenomenon could be explained by the fact that preformation of TFs in the cytoplasm may result in a faster response than *de novo* synthesis (Marathe et al., 2024). Our previous studies revealed that low-concentration CuSO_4_ treatment rapidly increased ethylene production by increasing *ACS8* transcription in *Arabidopsis* (Zhang et al., 2018). In *Solanum tuberosum*, the ET content increased to the peak level at 15 min after CuSO_4_ treatment (Liu et al., 2020). In this study, we found that bHLH107 was almost unable to enhance *Arabidopsis* disease resistance through transcriptional upregulation (Fig. 3A, 3B), whereas protein modifications may be induced faster than *de novo* synthesis. The transfer of bHLH107 from the cytoplasm to the nucleus is somewhat mediated via its amino acid (AA) residues 62 and 72, which are phosphorylated by CPK3 (Figs. 5A, 5B, 6A, 6B). Previous studies have shown that R/KXXS/T (X represents any amino acid) is a conserved motif phosphorylated by CDPKs (Furihata et al., 2006). Additionally, CPK3 phosphorylates several TFs, such as ERF1, HsfB2a and CZF1/ZFAR1, which are associated with the transcriptional activation of *DR* genes (Kanchiswamy et al., 2010). Here, we found that the conserved site of Ser72 in the 69-72 (KLLS) motif of bHLH107 can also be phosphorylated by CPK3 (Figs. 5B, S6). In addition to revealing its function in regulating the Cu^2+^-triggered immune response, we also revealed the phosphorylation mechanism for bHLH107. Previously, we reported that treatment with the calcium inhibitor LaCl_3_ could suppress Cu^2+^-induced resistance to *Pst* DC3000 (Liu et al., 2015). Here, we found that the Ca^2+^ sensor of CPK3 is directly involved in Cu^2+^-triggered *ACS8* expression and resistance. Therefore, our results revealed the existence of crosstalk between Ca^2+^ and Cu^2+^ signaling.

CPK3, as a Ca^2+^ sensor, is located not only in the cytosol and nucleus but also in guard cells to regulate ABA-activated plasma membrane Ca^2+^-permeable channels (Mori et al., 2006). Moreover, CPK3 regulates stomatal closure induced by flag22 and *Pst* DC3000 pathogenicity (Lu et al., 2020). In our previous studies, CuRE-35S mini:GFP was found to be highly expressed in guard cells, and Cu^2+^ promoted the reopening of stomata in leaves and induced the defense response to *Phytophthora infestans* by suppressing ABA synthesis (Zhang et al., 2018; Liu et al., 2020). The colocalization of CuRE-35S mini:GFP and CPK3 in guard cells provides additional possibilities for clustering the immune signals induced by Cu^2+^ and Ca^2+^. In *Arabidopsis*, CPKs phosphorylate 14-3-3 protein (Mehlmer et al., 2010; Swatek et al., 2014; Ormancey et al., 2017), resulting in regulation of downstream signaling pathways (Lachaud et al., 2013). The 14-3-3 protein typically regulates the function of phosphorylated proteins via protein-protein interactions, such as by changing their subcellular localization or participating in signal transduction (Huang et al., 2018; Chevalier et al., 2009). Under hypoxic stress, CPK12 is rapidly activated, translocating from the cytoplasm to the nucleus, potentiating plant hypoxia sensing by phosphorylating ERF-VII transcription factors, and the 14-3-3κ protein serves as a negative modulator of CPK12 cytosolic-nuclear translocation (Fan et al., 2023a). Consistent with these findings, we found that the CPK3-interacting proteins GRF1, GRF2 and GRF5 could also be bound by bHLH107 via IP-MS (Mehlmer et al., 2010; Swatek et al., 2014; Supplementary Table S2), indicating that 14-3-3 proteins may function as “immune response brakes” to fine tune Cu^2+^-induced resistance; this hypothesis needs to be further investigated.

In addition to being a coactivator of bHLH107 to directly regulate TF activity, HY5, as a transcriptional activator of light signaling, binds to 3894 genes (Lee et al., 2007), 38% of which are bound by GCN5, the histone acetyltransferase (Benhamed et al., 2008). We retrieved supplementary data on target genes bound by GCN5 or both GCN5 and HY5 published by Benhamed et al. in 2008; unfortunately, the *ACS8* gene was excluded, which indicated that Cu^2+^ may synergistically induce other acetyltransferases with HY5 to regulate *ACS8* gene transcription. Therefore, there are similar and differential characteristics of the expression patterns of target genes between light signaling and copper signaling. Although an increasing number of studies have reported that HY5 is required for histone acetylation of target genes induced by light signals (Guo et al., 2008; Charron et al., 2009; Velanis et al., 2016; Benhamed et al., 2006; Benhamed et al., 2008; Bertrand et al., 2005; Schenke et al., 2014a; Schenke et al., 2014b), an acetyltransferase that physically interacts with HY5 has not yet been identified. However, HY5 may mediate acetylation of the *ACS8* promoter to promote the bHLH107-induced *ACS8* expression upon Cu^2+^ treatment, which needs to be investigated further. Moreover, previous studies have shown that HY5 interacts with SPL7 in response to copper deficiency, thereby regulating the Cu^2+^ level in chloroplasts (Zhang et al., 2014). Moreover, HY5 inhibits the proliferation of downy mildew in *Arabidopsis* by promoting the upregulation of disease resistance-related genes and inducing ROS bursts (Chen et al., 2021). These findings suggest that HY5 participates not only in the regulation of copper homeostasis but also in the Cu^2+^-induced immune response in *Arabidopsis*.

In summary, our findings provide evidence that the CPK3-bHLH107-HY5-*ACS8* module regulates the conversion of Cu^2+^ signals into induction of ET production to modulate defense responses in *Arabidopsis* (Fig. 9). This study provides insight into the Cu^2+^-triggered immune response upstream of *ACS8*. More importantly, we revealed the transcriptional role and identified the novel phosphorylated target of CPK3, bHLH107, involved in the regulation of *ACS8* expression. In addition, HY5 cooperates with bHLH107 and binds to the G-box and CuRE respectively to trigger downstream immune responses in the nucleus (Fig 9).

## MATERIALS AND METHODS

### Plant materials and growth conditions

The wild-type *Arabidopsis* plants used in this study were of the Columbia (Col-0) and WS-0 ecotypes. The mutants were obtained from TAIR-ABRC: *bhlh107-1* (GABI_291G03), *bhlh107-2* (GABI_267B09), *acs8* (SALK_006628) (Zhang et al., 2018), *cpk3* (SALK_107620C), moreover, *ks50* (Oyama et al. 1997), and 35S::*HY5-GFP* (Shi et al., 2024) transgenic plants were generously provided by Professor Gang Li. The *bhlh107* line was generated in this study via CRISPR technology (Xie et al., 2015). The primers used for PCR-based genotyping of the mutant and transgenic plants are listed in Supplementary Table 4. *Arabidopsis* seeds were sown on half-strength Murashige and Skoog (MS) medium (Coolaber, Beijing, China) supplemented with 1.5% (w/v) sucrose and 0.8% (w/v) agar (pH 5.8) after surface sterilization. After stratification for 48 h at 4°C in the dark, the plates were incubated under a 16 h light/8 h dark photoperiod (20°C-22°C); 10-day-old seedlings were subsequently transplanted to soil for further growth in a greenhouse.

### Vector construction and generation of transgenic *Arabidopsis* plants

To generate a luciferase reporter driven by CuRE-35S mini, a CuRE with the 35S mini sequence was synthesized and inserted into the pGreen-0800-*Bam*H I-cut vector. To construct the *bHLH107*-HA vector, the *bHLH107* CDS was amplified from Col-0 cDNA via PCR and then inserted into the PS1300TB-HA-*Eco*RI-*Sac*I-cut (containing the 35S promoter and the sequence encoding an HA tag) vector. To generate the MBP-tagged *bHLH107* construct, the *bHLH107* CDS fragments were amplified and inserted into *Bam*H I-*Hin*dIII-digested pMAL-c2X. HIS-*HY5* and HIS-*CPK3* were cloned and inserted into *Bam*H I-*Hin*dIII- and *Bam*HI-*Sac* I-digested pET-28a, respectively. GST-*CPK3* was cloned and inserted into pGEX-4T1-*Eco*RI-*Sal* I-cut. The *HY5*-nLuc, *CPK3*-nLuc and *Actin*-nLuc (LOC_Os01g05490) genes were cloned as the respective DNA fragments and inserted into 35S::nLuc; cLuc-*bHLH107*, and cLuc-*Actin* was inserted into the 35S::cLuc vector. The genomic DNA of *bHLH107* with its 2-kb promoter and *CPK3* CDS fragment was inserted into P7A and PS1300T-MYC (containing the 35S promoter and the sequence encoding a MYC tag) vectors to generate the YFP-tagged bHLH107 and 35S::*CPK3* constructs, respectively. The genomic DNA of *bHLH107* and *ACS8* carrying each promoter was cloned and inserted into PC1300TB-Flag and pCAMBIA1300-mCherry to generate *ProbHLH107:gbHLH107* and *Pro_ACS8_:gACS8*, respectively. For the Y1H assay, *Kpn*I and *Sal*I sites were used to insert the various sequences of *ACS8* into the pLacZi2μ vector. The *HY5* CDS was ligated into the *Eco*RI-*Xho*I sites of the GAD vector (Lin et al., 2007). All primers used to generate the abovementioned constructs are listed in Supplementary Table 4.

### Bacterial growth assays

Bacterial growth assays were carried out as previously described (Zhang et al., 2018; Yu et al., 2021). *Pst* DC3000 was cultured in King’s B liquid medium for 36 h at 28°C, centrifuged at 6000 rpm for 5 min, and rinsed with 10 mM MgCl_2_. *Pst* DC3000 was infiltrated into the back of leaves. At 0 d and 3 d post-inoculation (dpi), the leaves were harvested, weighed, and surface sterilized. Afterward, the leaves were ground with 100 μl of sterile ddH_2_O and subsequently serially diluted (1:10) for plating on King’s B medium at 28°C for colony counting.

### Luciferase assays

To investigate bHLH107 binding to the CuRE in the *ACS8* promoter and protein-protein interactions, we inoculated HY5-nLuc-, CPK3-nLuc-, bHLH107-cLuc-, Actin-nLuc-, cLuc-Actin-, PS1300TB-HA-, 35Smini:Luc-, 35S::bHLH107-HA- and CuRE-35S mini:Luc-carrying Agrobacterium into Luria‒Bertani (LB) medium and incubated the cultures with shaking at 28°C for 2‒3 d. The mixture was subsequently centrifuged at 6000 rpm for 15 min and rinsed with an infection solution (10 mM MgCl_2_, 10 mM MES, 100 μM acetosyringone, 15 g/L sucrose, pH 5.7), the OD600 was adjusted to 1.0, and the bacteria were mixed in equal volumes. The mixture was incubated at 28°C for 2 h for infiltration into *N. benthamiana* with a needleless syringe, and the plants were grown in the greenhouse for 60-72 h. Luciferase signal detection was performed, and a luciferase substrate solution (100 mM Tris-HCl, pH 7.8; 150 μM ATP; 10 mM MgCl_2_; 3 g/L D-luciferin potassium salt) was prepared, which was sprayed on the backs of the leaves; the leaves were then incubated in darkness for 10 min. Fluorescence signals were detected via a NightSHADE LB985 plant living molecular imaging system (Berthold, Germany).

### EMSA

EMSAs were performed according to previously reported methods (Liu et al., 2020). The biotin-labeled probes listed in Supplementary Table 4 and a Chemiluminescent EMSA Kit (Beyotime, Shanghai, China) were used. Briefly, 2 μg of purified protein was incubated together with unlabeled or biotin-labeled wild-type or mutant probes. The binding reactions were incubated at 25°C for 20 min, and then the products were separated on 6% native polyacrylamide gels in TBE buffer. The labeled probes were detected according to the manufacturer’s instructions.

### ChIP‒qPCR

ChIP assays were performed as described previously, with slight modifications (Zhao et al., 2020). The samples were ground well, and 0.5 g of powder was resuspended in 1 mL of buffer F (50 mM HEPES-KOH pH 7.5, 150 mM NaCl, 1 mM EDTA, 1% Triton X-100, 0.1% sodium deoxycholate, 1× protein inhibitor cocktail). After crosslinking, the mixture was quenched with 0.125 M glycine. The mixture was centrifuged at 12000 × g, and then the pellet was washed. The pellet was resuspended in 1 mL of buffer S (50 mM HEPES-KOH pH 7.5, 150 mM NaCl, 1 mM EDTA, 1% Triton X-100, 0.1% sodium deoxycholate, 1% SDS, protease inhibitor) in a 1.5 ml tube. After sonication, the sample was centrifuged at 12000 × g for 5 min, and 20 µL of the supernatant was used as the input sample, whereas the remaining supernatant was diluted 5 times with buffer F for binding with antibodies overnight (binding with Protein A/G magnetic beads (HY-K0202, MedChemExpress) for approximately 4 h before use). The beads were subsequently washed twice with low-salt buffer (50 mM HEPES-KOH pH 7.5, 150 mM NaCl, 1% Triton X-100, 1 mM EDTA, 0.1% sodium deoxycholate, 0.1% SDS), twice/three times with high-salt buffer (50 mM HEPES-KOH pH 7.5, 500 mM NaCl, 1% Triton X-100, 1 mM EDTA, 0.1% sodium deoxycholate, 0.1% SDS), twice with LiCl buffer (10 mM Tris-HCl 8.0, 250 mM LiCl, 0.5% CA-630, 0.1% sodium deoxycholate, 1 mM EDTA), and twice with TE buffer (10 mM Tris-HCl pH 8.0, 1 mM EDTA). The beads were resuspended in 100 µL of elution buffer (50 mM Tris-HCl pH 7.5, 10 mM EDTA, and 1% SDS). The DNA was purified via a phase lock gel (Tiangen, Beijing, China). The precipitated DNA samples were quantified by qPCR using the respective pair of primers listed in Supplementary Table 4.

### Yeast one-hybrid assays

For yeast one-hybrid assays assays, full-length *HY5* and various *LacZ* reporter plasmids were cotransformed into EGY48 competent yeast cells, which were subsequently plated on SD medium lacking Trp and Ura (SD/-Trp-Ura, Coolaber, Beijing, China), and positive clones were transferred to SD/-Trp-Ura medium that contained BU salts, galactose, raffinose, and X-gal (Lin et al., 2007).

### Coimmunoprecipitation analysis

Co-IP analysis was performed according to previously reported methods (Yang et al., 2023b). Briefly, total protein was extracted from tobacco leaves with co-IP buffer (50 mM Tris HCl pH 7.5; 150 mM NaCl; 5 mM MgCl_2_; 5% glycerol; 0.1% NP-40; 1 mM DTT; 1 mM PMSF; 1× protein inhibitor cocktail). The protein extract was incubated with magnetic bead-conjugated mouse anti-DDDDK-Tag monoclonal antibody (mAb) (ABclone, Wuhan, China) for 2 h. After incubation, the beads were washed four times with wash buffer (50 mM Tris HCl pH 7.5; 300 mM NaCl; 5 mM MgCl_2_; 5% glycerol; 0.1% NP-40; 1 mM DTT; 1 mM PMSF; 1x protein inhibitor cocktail), and the samples were subjected to western blot analysis via an anti-HA tag mouse mAb and an anti-Flag tag mouse mAb (Proteintech, Wuhan, China).

### *In vitro* pull-down assays

*In vitro* pull-down assays were performed according to previously reported methods (Wu et al., 2022). Proteins with different tags were expressed in *E.* coli (BL21 DE3) cells. For *in vitro* pull-down assays, bait proteins (MBP, MBP-bHLH107, GST, and GST-CPK3) were mixed with each prey protein in PBS buffer with a 1 × protein inhibitor cocktail. After incubation at 4°C for 1 h, different tagged beads relevant to tags for bait proteins were added and incubated for another hour. Then, the beads were washed six times with PBS, and the proteins that bound to the beads were eluted with elution buffer (50 mM Tris-HCl pH 8.0, 1 mM EDTA, 10 mM DTT, 0.1% Triton X-100, 10 mM maltose or 10 mM GSH) at room temperature for 10 min. In the next step, the proteins were denatured with 5 × sodium dodecyl sulfate (SDS) loading buffer at 95°C for 10 min. The input and eluted proteins were separated on 10% sodium dodecyl sulfate‒ polyacrylamide gel electrophoresis (SDS‒PAGE) gels and immunoblotted with an anti-MBP tag mouse mAb and an anti-GST tag mouse mAb (Proteintech, Wuhan, China), respectively.

### RNA extraction and RT‒qPCR

Total RNA was extracted from 4-week-old *Arabidopsis* plants via a Plant RNA Kit (OMEGA Biotek, Norcross, USA). cDNA was synthesized via ReverTra Ace® qPCR RT Master Mix with gDNA Remover (TOYOBO). RT‒qPCR was performed via ChamQ SYBR qPCR Master Mix (without ROX) (Vazyme, Jiangsu, China) on a CFX Connect^TM^ Real-Time System and a CFX384^TM^ Real-Time System (Bio-Rad). PCRs were performed in triplicate or quadruplicate for each sample, and the relative expression levels were normalized to that of *Actin 2*. All the qRT‒PCR primers used are listed in Supplementary Table 4. Materials for RT‒qPCR assays were collected from two or three replicates.

### Kinase assay

One microgram of MBP-bHLH107 protein was incubated with equal amounts of GST-CPK3 protein in kinase assay buffer (50 mM Tris-HCl pH 7.5, 1 mM CaCl_2_, 10 mM MgCl_2_, 1 mM DTT, 250 μM ATP) at 30°C for 1 h and 2 h. The reactions were then mixed with 5×SDS loading buffer and incubated at 95°C for 10 min. The proteins were separated on 10% SDS‒PAGE gels and 8% Phos-tag gels (Vazyme, Nanjing, China) and immunoblotted with an anti-MBP tag mouse mAb and an anti-GST tag mouse mAb (Proteintech, Wuhan, China), respectively.

### Protoplast transformation

The specific experimental steps followed a previously described protocol (Li et al., 2008), with slight modifications. In brief, 21-d-old *Arabidopsis* leaves were cut into thin filaments and placed in an enzymatic mixture (0.4 M mannitol, 20 mM KCl, 20 mM MES-KOH pH 5.7, cellulase R10, macerozyme R10, 10 mM CaCl_2_, 5 mM β-mercaptoethanol, 0.1% FBS). The mixture was incubated at room temperature for 90 min with shaking at 50 rpm. After filtering through a cell sieve, the mixture was centrifuged at 100 × g for 3 min, resuspended in 10 ml of W5 solution (154 mM NaCl, 125 mM CaCl_2_, 5 mM KCl, 2 mM MES-KOH pH 5.7), incubated on ice for 40 min, centrifuged at 100 × g for 3 min, and resuspended in MMG solution (0.4 M mannitol, 15 mM MgCl_2_, 4 mM MES-KOH pH 5.7). A total of 10 μg of plasmid (1 μg/μl), 200 μl of protoplasts, and 220 μl of transformation solution (40% PEG 4000, 200 mM mannitol, 100 mM CaCl_2_) were mixed in a 2 ml centrifuge tube, incubated at room temperature for 25 min, resuspended in 800 μl of W5 solution to terminate the reaction, centrifuged at 100 × g for 3 min, resuspended in 2 ml of W5 solution, and incubated in the dark for 16 h. Finally, the fluorescence signal was detected via a Microsystems confocal microscope (Leica, Wetzlar, Germany) with a 20 × objective. The following Leica filter sets were used in this study: EYFP (514 nm) and mCherry (552 nm).

### Protein extraction

For nuclear-cytosolic fractionation, approximately 0.3 g of leaves were ground to a fine powder in liquid nitrogen, and 3 mL of lysis buffer (20 mM Tris-HCl pH 7.5, 20 mM KCl, 2 mM EDTA, 2.5 mM MgCl_2_, 25% glycerol, 250 mM sucrose, 5 mM DTT, 0.4 mM PMSF, 1 × protein inhibitor cocktail) was added to the powder. The homogenate was filtered through a double layer of Miracloth. The flow-through sample is referred to here as “Total”. The flow-through sample was spun at 1500 × g for 10 min. The supernatant, consisting of the cytoplasmic fraction, was centrifuged at 10,000 × g for 10 min at 4°C, and the resulting supernatant was collected and designated “Cytosol”. The pellet was subsequently washed four times with 1 mL of NRBT nuclear resuspension buffer (20 mM Tris-HCl pH 7.4, 25% glycerol, 2.5 mM MgCl_2_, 0.2% Triton X-100, 0.4 mM PMSF, protease inhibitor cocktail). The washed nuclei were resuspended in 500 μL of NRB2 buffer (20 mM Tris-HCl pH 7.5, 0.25 M sucrose, 10 mM MgCl_2_, 0.5% Triton X-100, 5 mM β-mercaptoethanol, 0.4 mM PMSF, 1 × protein inhibitor cocktail). Nuclei were carefully overlaid on top of 500 μL of NRB3 (20 mM Tris-HCl pH 7.5, 1.7 M sucrose, 10 mM MgCl_2_, 0.5% Triton X-100, 5 mM β-mercaptoethanol, 0.4 mM PMSF, protease inhibitor cocktail) and centrifuged at 16,000 × g for 45 min at 4°C, and the pellets were collected and designated “nuclei”. The nuclei were eluted with nuclear lysis buffer (50 mM Tris-HCl pH 8.0, 10 mM EDTA, 1% SDS). The samples were boiled at 95°C for 10 min and then subjected to western blot analysis.

### DNA pull-down MS

Nuclear protein was extracted from 20 g of 4-week-old Col-0 plants after they were sprayed with 100 μM CuSO_4_ (0.02% Tween-20) via a CelLytic™ PN kit (Sigma, USA). The CuRE with/without biotin labeling was synthesized by Sangon (Shanghai, China). The CuRE was incubated with Dynabeads™ Streptavidin for target enrichment (Thermo Fisher Scientific, USA) at 4°C for 3 h. Afterward, the nuclear protein suspension was incubated with CuRE-bound Dynabeads at 4°C for 3 h, followed by enrichment and washing five times with wash solution (50 mM Tris HCl pH 7.5; 150 mM NaCl; 1 mM EDTA; 0.1% SDS; 1% Triton X-100; 5 mM DTT; 1 mM PMSF; 1x protein inhibitor cocktail) at 4°C to remove nonspecifically bound proteins. Eighty microliters of 1x SDS loading buffer was added to the beads, which were subsequently incubated at room temperature for 10 min for denaturation at 95°C for 10 min, after which the samples were subjected to SDS‒PAGE and Coomassie brilliant blue staining. The gel strips were cut out for mass spectrometry detection (Applied Protein Technology, Shanghai, China) to decode the peptide information for bound proteins.

### IP‒MS

Total protein was extracted from 20 g of 4-week-old 35S::*bHLH107*-*HA*/Col-0 transgenic seedlings after they were sprayed with 100 μM CuSO_4_ (0.02% Tween-20) or not. After incubation with anti-HA magnetic beads (MedChemExpress, USA) for 3 hours, 80 μl of 1x SDS loading buffer was added, and the mixture was incubated at room temperature for 10 minutes and denatured at 95°C for 10 minutes. The samples were then subjected to SDS‒PAGE and Coomassie Brilliant Blue staining. The gel strips were cut out for mass spectrometry detection (Applied Protein Technology, Shanghai, China) to decode the peptide information for bound proteins.

## ACKNOWLEDGMENTS

We thank Prof. Gang Li from College of Life Sciences, Shandong Agricultural University for providing the seeds of HY5 overexpressing lines and *hy5* mutant.

## AUTHORS CONTRIBUTIONS

Z.C. and H.L. designed the experiments. Y.Y. performed the experiments and analyzed data. Z.C. and H.L. conceived and supervised the project. Y.Y., Z.C. and X.C. wrote the manuscript.

---

**Table S1** List of proteins pull-down by CuRE *cis*-element upon CuSO_4_ treatment.

**Table S2** List of proteins IP by bHLH107 upon CuSO_4_ treatment or not.

**Table S3** List of phosphorylated-peptides of bHLH107 phosphorylated by CPK3.

**Table S4** Primers used in this study.

## Reference

Behlau F, Hong JC, Jones JB, Graham JH. Evidence for acquisition of copper resistance genes fro different sources in citrus-associated xanthomonads. Phytopathology 2013: 103(5): 409–418. 10.1094/PHYTO-06-12-0134-R

Bender KW, Zielinski RE, Huber SC. Revisiting paradigms of Ca^2+^ signaling protein kinase regulation in plants. Biochem. J. 2018: 475(1): 207–223. 10.1042/BCJ20170022

Benhamed M, Bertrand C, Servet C, Zhou DX. *Arabidopsis GCN5*, *HD1*, and *TAF1/HAF2* interact to regulate histone acetylation required for light-responsive gene expression. Plant Cell 2006: 18(11): 2893–2903. 10.1105/tpc.106.043489

Benhamed M, Martin-Magniette ML, Taconnat L, Bitton F, Servet C, De Clercq R, De Meyer B, Buysschaert C, Rombauts S, Villarroel R, et al. Genome-scale Arabidopsis promoter array identifies targets of the histone acetyltransferase GCN5. Plant J. 2008:56(3): 493–504. 10.1111/j.1365-313X.2008.03606.x

Bertrand C, Benhamed M, Li YF, Ayadi M, Lemonnier G, Renou JP, Delarue M, Zhou DX. *Arabidopsis* HAF2 gene encoding TATA-binding protein (TBP)-associated factor TAF1, is required to integrate light signals to regulate gene expression and growth. J. Biol. Chem. 2005: 280(2): 1465–1473.

Burkhead JL, Gogolin Reynolds KA, Abdel-Ghany SE, Cohu CM, Pilon M. Copper homeostasis. New Phytol. 2009: 182(4):799-816. 10.1111/j.1469-8137.2009.02846.x

Carretero-Paulet L, Galstyan A, Roig-Villanova I, Martínez-García JF, Bilbao-Castro JR, Robertson DL. Genome-wide classification and evolutionary analysis of the bHLH family of transcription factors in Arabidopsis, poplar, rice, moss, and algae. Plant Physiol. 2010: 153(3): 1398–1412. 10.1104/pp.110.153593

Charron JB, He H, Elling AA, Deng XW. Dynamic landscapes of four histone modifications during deetiolation in *Arabidopsis*. Plant Cell 2009: 21(12): 3732–3748. 10.1105/tpc.109.066845

Chattopadhyay S, Ang LH, Puente P, Deng XW, Wei N. Arabidopsis bZIP protein HY5 directly interacts with light-responsive promoters in mediating light control of gene expression. Plant Cell 1998: 10(5): 673–683. 10.1105/tpc.10.5.673

Chen JN, Wu LT, Song K, Zhu YS, Ding W. Nonphytotoxic copper oxide nanoparticles are powerful “nanoweapons” that trigger resistance in tobacco against the soil-borne fungal pathogen *Phytophthora nicotianae*. J. Integr. Agr. 2022a: 21(11). 10.1016/j.jia.2022.08.086.

Chen QB, Wang WJ, Zhang Y, Zhan QD, Liu K., Botella JR, Bai L, Song CP. Abscisic acid-induced cytoplasmic translocation of constitutive photomorphogenic 1 enhances reactive oxygen species accumulation through the HY5-ABI5 pathway to modulate seed germination. Plant Cell Environ. 2022b: 45(5): 1474–1489. 10.1111/pce.14298

Chen SY, Ma T, Song SR, Li XL, Fu P, Wu W, Liu JQ, Gao Y, Ye WX, Dry IB, et al. Arabidopsis downy mildew effector HaRxLL470 suppresses plant immunity by attenuating the DNA-binding activity of bZIP transcription factor HY5. New Phytol. 2021: 230(4): 1562–1577. 10.1111/nph.17280

Cheng HK, Pan GY, Zhou N, Zhai ZK, Yang LQ, Zhu HF, Cui X, Zhao PY, Zhang HF, Li SJ, et al. Calcium-dependent Protein Kinase 5 (CPK5) positively modulates drought tolerance through phosphorylating ABA-Responsive Element Binding Factors in oilseed rape (*Brassica* napus L.). Plant Sci. 2022: 315: 111125. 10.1016/j.plantsci.2021.111125

Chevalier D, Morris ER, Walker JC. 14-3-3 and FHA domains mediate phosphoprotein interactions. Annu Rev. Plant Biol. 2009: 60: 67–91. 10.1146/annurev.arplant.59.032607.092844

Chu LT, Yang C, Zhuang F, Gao YM, Luo M. The HDA9-HY5 module epigenetically regulates flowering time in *Arabidopsis thaliana*. J. Cell Physiol. 2022: 237(7): 2961–2968. 10.1002/jcp.30761

Cui X, Zhao PY, Liang WW, Cheng, Q, Mu BB, Niu FF, Yan JL, Liu CL, Xie H, Kav NNV, et al. A Rapeseed WRKY Transcription Factor Phosphorylated by CPK Modulates Cell Death and Leaf Senescence by Regulating the Expression of ROS and SA-Synthesis-Related Genes. J. Agr. Food Chem. 2020: 68(28): 7348–7359. 10.1021/acs.jafc.0c02500

Drepper F, Hippler M, Nitschke W, Haehnel W. Binding dynamics and electron transfer between plastocyanin and photosystem I. Biochemistry-Us 1996: 35(4): 1282–1295. 10.1021/bi951471e

Fan B, Liao K, Wang LN, Shi LL, Zhang Y, Xu LJ, Zhou Y, Li JF, Chen YQ, Chen QF, et al. Calcium-dependent activation of CPK12 facilitates its cytoplasm-to-nucleus translocation to potentiate plant hypoxia sensing by phosphorylating ERF-VII transcription factors. Mol. Plant 2023a: 16(6): 979–998. 10.1016/j.molp.2023.04.002

Fan W, Liao XL, Tan YQ, Wang XR, Schroeder JI, Li ZX. Arabidopsis PLANT U-BOX44 down-regulates osmotic stress signaling by mediating Ca^2+^-DEPENDENT PROTEIN KINASE4 degradation. Plant Cell 2023b: 35(10): 3870–3888. 10.1093/plcell/koad173

Furihata T, Maruyama K, Fujita Y, Umezawa T, Yoshida R, Shinozaki K, Yamaguchi-Shinozaki K. Abscisic acid-dependent multisite phosphorylation regulates the activity of a transcription activator AREB1. P. Natl. Acad. Sci. Usa. 2006: 103(6): 1988–1993. 10.1073/pnas.0505667103

Gao F, Dubos, C. The arabidopsis bHLH transcription factor family. Trends in plant Plant sci. 2023: S1360–1385(23)00381-3. Advance online publication. 10.1016/j.tplants.2023.11.022

Gong W, He K, Covington M, Dinesh-Kumar SP, Snyder M, Harmer SL, Zhu YX, Deng XW. The development of protein microarrays and their applications in DNA-protein and protein-protein interaction analyses of *Arabidopsis* transcription factors. Mol. Plant 2008: 1(1): 27–41. 10.1093/mp/ssm009

Gou Y, Jing Y, Song J, Nagdy MM, Peng C, Zeng L, Chen M, Lan X, Htun ZLL, Liao Z, et al. A novel bHLH gene responsive to low nitrogen positively regulates the biosynthesis of medicinal tropane alkaloids in *Atropa belladonna*. Int. J. Biol. Macromol. 2024: 266(Pt 1): 131012. Advance online publication. 10.1016/j.ijbiomac.2024.131012

Guo F, Meng XQ, Hong HT, Liu SY, Yu J, Huang C, Dong TT, Geng HX, Li ZY, Zhu MK. Systematic identification and expression analysis of *bHLH* gene family reveal their relevance to abiotic stress response and anthocyanin biosynthesis in sweetpotato. BMC Plant Biol. 2024a: 24(1): 156. 10.1186/s12870-024-04788-0

Guo HY, Xu CT, Wang F, Jiang LQ, Zhang YH, Wang LF, Liu DY, Zhao JC, Xia C, Gu Y, et al. Transcriptome analysis and functional verification reveal the roles of copper in resistance to potato virus Y infection in tobacco. Pestic. Biochem. Phys. 2024b: 201: 105893. 10.1016/j.pestbp.2024.105893

Guo L, Zhou J, Elling AA, Charron JB, Deng XW. Histone modifications and expression of light-regulated genes in Arabidopsis are cooperatively influenced by changing light conditions. Plant Physiol. 2008: 147(4): 2070–2083. 10.1104/pp.108.122929

Harper JF., Breton G, Harmon A. Decoding Ca^2+^ signals through plant protein kinases. Annu. Rev. Plant Biol. 2004: 55: 263–288. 10.1146/annurev.arplant.55.031903.141627

Hu YF, Zhao HY, Xue LY, Nie N, Zhang H, Zhao N, He SZ, Liu QC, Gao SP, Zhai H. *IbMYC2* Contributes to Salt and Drought Stress Tolerance via Modulating Anthocyanin Accumulation and ROS-Scavenging System in Sweet Potato. Int. J. Mol. Sci. 2024: 25(4): 2096. 10.3390/ijms25042096

Huang X, Zhang Q, Jiang YP, Yang CW, Wang QY, Li L. Shade-induced nuclear localization of PIF7 is regulated by phosphorylation and 14-3-3 proteins in Arabidopsis. Elife 2018: 7: e31636. 10.7554/eLife.31636

Jiang WB, Yin QG, Liu JY, Su XJ, Han XY, Li Q, Zhang J, Pang YZ. The APETALA2-MYBL2 module represses proanthocyanidin biosynthesis by affecting formation of the MBW complex in seeds of *Arabidopsis thaliana*. Plant Commun. 2024: 5(3): 100777. 10.1016/j.xplc.2023.100777

Kanchiswamy CN, Takahashi H, Quadro S, Maffei ME, Bossi S, Bertea C, Zebelo SA, Muroi A, Ishihama N, Yoshioka H, et al. Regulation of Arabidopsis defense responses against Spodoptera littoralis by CPK-mediated calcium signaling. Bmc Plant Biol. 2010: 10: 97. 10.1186/1471-2229-10-97

Kelly G, Yaaran A, Gal A, Egbaria A, Brandsma D, Belausov E, Wolf D, David-Schwartz R, Granot D, Eyal Y, et al. Guard cell activity of PIF4 and HY5 control transpiration. Plant Sci. 2023: 328: 111583. 10.1016/j.plantsci.2022.111583

Kim JG, Mudgett MB. Tomato bHLH132 Transcription Factor Controls Growth and Defense and Is Activated by *Xanthomonas euvesicatoria* Effector XopD During Pathogenesis. Mol. Plant Microbe. In. 2019: 32(12): 1614–1622. 10.1094/MPMI-05-19-0122-R

Kuras L, Barbey R, Thomas D. Assembly of a bZIP-bHLH transcription activation complex: formation of the yeast Cbf1-Met4-Met28 complex is regulated through Met28 stimulation of Cbf1 DNA binding. Embo J. 1997: 16(9): 2441–2451. 10.1093/emboj/16.9.2441

Lachaud C, Prigent E, Thuleau P, Grat S, Da Silva D, Brière C, Mazars C, Cotelle V. 14-3-3-regulated Ca^2+^-dependent protein kinase CPK3 is required for sphingolipid-induced cell death in *Arabidopsis*. Cell Death Differ. 2013: 20(2): 209–217. 10.1038/cdd.2012.114

Lamichhane JR, Osdaghi E, Behlau F, Köhl J, Jones J, Aubertot JN. Thirteen decades of antimicrobial copper compounds applied in agriculture. A review. Agron. Sustain. Dev. 2018: 38(28). 10.1007/s13593-018-0503-9

Lee J, He K, Stolc V, Lee H, Figueroa P, Gao Y, Tongprasit W, Zhao H, Lee I, Deng XW. Analysis of transcription factor HY5 genomic binding sites revealed its hierarchical role in light regulation of development. Plant Cell 2007: 19(3): 731–749. 10.1105/tpc.106.047688

Leivar P, Quail PH. PIFs: pivotal components in a cellular signaling hub. Trends Plant Sci. 2011: 16(1): 19–28. 10.1016/j.tplants.2010.08.003

Li XY, Chanroj S, Wu ZY, Romanowsky SM, Harper JF, Sze H. A distinct endosomal Ca^2+^/Mn^2+^ pump affects root growth through the secretory process. Plant Physiol. 2008: 147(4): 1675–1689.

Li ZY, Fu YJ, Wang YZ, Liang JS. Scaffold protein RACK1 regulates BR signaling by modulating the nuclear localization of BZR1. New Phytol. 2023: 239(5): 1804–1818. 10.1111/nph.19049

Libault M, Wan J, Czechowski T, Udvardi M, Stacey G. Identification of 118 *Arabidopsis* transcription factor and 30 ubiquitin-ligase genes responding to chitin, a plant-defense elicitor. Mol. Plant Microbe In. 2007: 20(8): 900–911. 10.1094/MPMI-20-8-0900

Lin RC, Ding L, Casola C, Ripoll DR, Feschotte C, Wang H. Transposase-derived transcription factors regulate light signaling in *Arabidopsis*. Science 2007: 318(5854): 1302-1305. 10.1126/science.1146281

Liu HF, Xue XJ, Yu Y, Xu MM, Lu CC, Meng XL, Zhang BG, Ding XH, Chu ZH. Copper ions suppress abscisic acid biosynthesis to enhance defence against Phytophthora infestans in potato. Mol. Plant Pathol. 2020: 21: 636–651.

Liu JZ, Shen YT, Cao HX, He K, Chu ZH, Li N. OsbHLH057 targets the AATCA cis-element to regulate disease resistance and drought tolerance in rice. Plant Cell Rep. 2022: 41(5): 1285–1299. 10.1007/s00299-022-02859-w

Liu YY, Zhang Q, Chen DX, Shi WS, Gao XM, Liu Y, Hu B, Wang AH, Li XY, An XY, et al. Positive regulation of ABA signaling by MdCPK4 interacting with and phosphorylating MdPYL2/12 in Arabidopsis. J. Plant Physiol. 2024: 293: 154165. 10.1016/j.jplph.2023.154165

Liu, HF, Zhang BG, Wu T, Ding Y, Ding XH, Chu ZH. Copper ion elicits defense response in *Arabidopsis* thaliana by activating salicylate- and ethylene-dependent signaling pathways. Mol. Plant 2015: 8: 1550–1553.

Lu YJ, Li P, Shimono M, Corrion A, Higaki T, He SY, Day B. Arabidopsis calcium-dependent protein kinase 3 regulates actin cytoskeleton organization and immunity. Nat. Commun. 2020: 11(1): 6234. 10.1038/s41467-020-20007-4

Ma XC, Sheng L, Li FF, Zhou T, Guo J, Chang YY, Yang J, Jin YF, Chen YW, Lu XP. Seasonal drought promotes citrate accumulation in citrus fruit through the CsABF3-activated CsAN1-CsPH8 pathway. New Phytol. 2024: 242(3): 1131–1145. 10.1111/nph.19671

Mankotia S, Jakhar P, Satbhai SB. HY5: a key regulator for light-mediated nutrient uptake and utilization by plants. New Phytol. 2024: 241(5): 1929–1935. 10.1111/nph.19516

Mankotia S, Singh D, Monika K, Kalra M, Meena H, Meena V, Yadav RK, Pandey AK, Satbhai SB. ELONGATED HYPOCOTYL 5 regulates BRUTUS and affects iron acquisition and homeostasis in *Arabidopsis thaliana*. Plant J. 2023: 114(6): 1267–1284. 10.1111/tpj.16191

Marathe S, Grotewold E, Otegui MS. Should I stay or should I go? Trafficking of plant extra-nuclear transcription factors. Plant Cell 2024: 36(5): 1524–1539. 10.1093/plcell/koad277

Massari ME, Murre C. Helix-loop-helix proteins: regulators of transcription in eucaryotic organisms. Mol. Cell Biol. 2000: 20(2): 429–440. 10.1128/MCB.20.2.429-440.2000

Mehlmer N, Wurzinger B, Stael S, Hofmann-Rodrigues D, Csaszar E, Pfister B, Bayer R, Teige M. The Ca^2+^ -dependent protein kinase CPK3 is required for MAPK-independent salt-stress acclimation in Arabidopsis. Plant Journal J. 2010: 63(3): 484–498.

Meng FW, Yang C, Cao JD, Chen H, Pang JH, Zhao QQ, Wang ZY, Qing Fu Z, Liu J. A bHLH transcription activator regulates defense signaling by nucleo-cytosolic trafficking in rice. J. Integr. Plant Biol. 2020: 62(10): 1552**-**1573. 10.1111/jipb.12922

Mir AR, Pichtel J, Hayat S. Copper: uptake, toxicity and tolerance in plants and management of Cu-contaminated soil. Biometals 2021: 34(4): 737–759. 10.1007/s10534-021-00306-z

Mori IC, Murata Y, Yang Y, Munemasa S, Wang YF, Andreoli S, Tiriac H, Alonso JM, Harper JF, Ecker JR, et al. CDPKs CPK6 and CPK3 function in ABA regulation of guard cell S-type anion- and Ca^2+^-permeable channels and stomatal closure. Plos Biol. 2006: 4(10): e327. 10.1371/journal.pbio.0040327

Onohata T, Gomi K. Overexpression of jasmonate-responsive *OsbHLH034* in rice results in the induction of bacterial blight resistance via an increase in lignin biosynthesis. Plant Cell Rep. 2020: 39(9): 1175–1184. 10.1007/s00299-020-02555-7

Ormancey M, Thuleau P, Mazars C, Cotelle V. CDPKs and 14-3-3 Proteins: Emerging Duo in Signaling. Trends Plant Sci. 2017: 22(3): 263–272.

Oyama T, Shimura Y, Okada K. The *Arabidopsis HY5* gene encodes a bZIP protein that regulates stimulus-induced development of root and hypocotyl. Gene. Dev. 1997: 11(22): 2983–2995. 10.1101/gad.11.22.2983

Pan GY, Zhang HF, Chen BY, Gao SD, Yang B, Jiang YQ. Rapeseed calcium-dependent protein kinase CPK6L modulates reactive oxygen species and cell death through interacting and phosphorylating RBOHD. Biochem. Bioph. Res. Co. 2019: 518(4): 719–725. 10.1016/j.bbrc.2019.08.118

Prodhan MY, Munemasa S, Nahar MN, Nakamura Y, Murata Y. Guard Cell Salicylic Acid Signaling Is Integrated into Abscisic Acid Signaling via the Ca^2+^/CPK-Dependent Pathway. Plant Physiol. 2018: 178(1): 441–450. 10.1104/pp.18.00321

Ranjan R, Malik N, Sharma S, Agarwal P, Kapoor S, Tyagi AK. OsCPK29 interacts with MADS68 to regulate pollen development in rice. Plant Sci. 2022: 321: 111297. 10.1016/j.plantsci.2022.111297

Rezayian M, Zarinkamar F. Nitric oxide, calmodulin and calcium protein kinase interactions in the response of *Brassica napus* to salinity stress. Plant Biology 2023: 25(3): 411–419. 10.1111/plb.13511

Schenke D, Cai D, Scheel D. Suppression of UV-B stress responses by flg22 is regulated at the chromatin level via histone modification. Plant Cell Environ. 2014a: 37(7): 1716–1721. 10.1111/pce.12283

Schenke D, Cai D. The interplay of transcription factors in suppression of UV-B induced flavonol accumulation by flg22. Plant Signal Behav. 2014b: 9(4): e28745. Advance online publication. 10.4161/psb.28745

Schulze S, Dubeaux G, Ceciliato PHO, Munemasa S, Nuhkat M, Yarmolinsky D, Aguilar J, Diaz R, Azoulay-Shemer T, Steinhorst L, et al. A role for calcium-dependent protein kinases in differential CO_2_ - and ABA-controlled stomatal closing and low CO_2_ -induced stomatal opening in Arabidopsis. New Phytol. 2021: 229(5): 2765–2779. 10.1111/nph.17079

Shi QB, Xia Y, Xue N, Wang QB, Tao Q, Li MJ, Xu D, Wang XF, Kong FY, Zhang HS, Li G. Modulation of starch synthesis in *Arabidopsis* via phytochrome B-mediated light signal transduction. J. Integr. Plant Biol. 2024: 66(5): 973-985. 10.1111/jipb.13630

Soriano GM, Ponamarev MV, Tae GS, Cramer WA. Effect of the interdomain basic region of cytochrome f on its redox reactions in vivo. Biochemistry-Us 1996: 35(46): 14590–14598. 10.1021/bi9616211

Sun YX, Li Q, Wu MD, Wang QW, Zhang DP, Gao Y. Rice PIFs: Critical regulators in rice development and stress response. Plant Mol Biol. 2024: 114(1): 1. 10.1007/s11103-023-01406-9

Swatek KN, Wilson RS, Ahsan N, Tritz RL, Thelen JJ. Multisite phosphorylation of 14-3-3 proteins by calcium-dependent protein kinases. Biochemical Journal J. 2014: 459(1): 15–25.

Thorstensen T, Grini PE, Mercy IS, Alm V, Erdal S, Aasland R, Aalen, RB. The Arabidopsis SET-domain protein ASHR3 is involved in stamen development and interacts with the bHLH transcription factor ABORTED MICROSPORES (AMS). Plant Mol. Biol. 2008: 66(1-2): 47–59. 10.1007/s11103-007-9251-y

Tian HZ, Fan GL, Xiong XW, Wang H, Zhang SQ, Geng GD. Characterization and transformation of the *CabHLH18* gene from hot pepper to enhance waterlogging tolerance. Front Plant Sci. 2024: 14: 1285198. 10.3389/fpls.2023.1285198

Toledo-Ortiz G, Huq E, Quail PH. The Arabidopsis basic/helix-loop-helix transcription factor family. Plant Cell 2003: 15(8): 1749–1770. 10.1105/tpc.013839

Trofimov K, Gratz R, Ivanov R, Stahl Y, Bauer, Brumbarova T. FER-like iron deficiency-induced transcription factor (FIT) accumulates in nuclear condensates. J. Cell Biol. 2024: 223(4): e202311048. 10.1083/jcb.202311048

Varympopi A, Dimopoulou A, Papafotis D, Avramidis P, Sarris I, Karamanidou T, Kerou, AK, Vlachou A, Vellis E, Giannopoulos A, et al. Antibacterial Activity of Copper Nanoparticles against *Xanthomonas campestris* pv. *vesicatoria* in Tomato Plants. Int. J. Mol. Sci. 2022: 23(8): 4080. 10.3390/ijms23084080

Velanis CN, Herzyk P, Jenkins GI. Regulation of transcription by the Arabidopsis UVR8 photoreceptor involves a specific histone modification. Plant Mol. Biol. 2016: 92(4-5): 425–443. 10.1007/s11103-016-0522-3

Voloudakis AE, Reignier TM, Cooksey DA. Regulation of resistance to copper in *Xanthomonas axonopodis* pv. *vesicatoria*. Appl. Environ. Microb. 2005: 71(2): 782–789. 10.1128/AEM.71.2.782-789.2005

Wang JJ, Gao J, Li W, Liu JX. CCaP1/CCaP2/CCaP3 Interact with Plasma Membrane H^+^-ATPases and Promote Thermo-Responsive Growth through Regulating Cell Wall Modification in *Arabidopsis*. Plant Commun. 2024: 100880. Advance online publication. 10.1016/j.xplc.2024.100880

Wang R, Yu MM, Xia JQ, Xing JP, Fan XP, Xu QH, Cang J, Zhang D. Overexpression of *TaMYC2* confers freeze tolerance by ICE-CBF-COR module in *Arabidopsis thaliana*. Front. Plant Sci. 2022: 13: 1042889. 10.3389/fpls.2022.1042889

Wang XN, Lv S, Han XY, Guan XJ, Shi X, Kang JK, Zhang LS, Cao B, Li C, Zhang W, et al. The Calcium-Dependent Protein Kinase CPK33 Mediates Strigolactone-Induced Stomatal Closure in *Arabidopsis thaliana*. Front Plant Sci. 2019: 10: 1630. 10.3389/fpls.2019.01630

Wu T, Zhang HM, Yuan B, Liu HF, Kong LG, Chu ZH, Ding XH. Tal2b targets and activates the expression of OsF3H_03g_ to hijack OsUGT74H4 and synergistically interfere with rice immunity. New Phytol. 2022: 233(4): 1864–1880. 10.1111/nph.17877

Xie KB, Minkenberg B, Yang YN. Boosting CRISPR/Cas9 multiplex editing capability with the endogenous tRNA-processing system. P. Natl. Acad. Sci. USA. 2015: 112(11): 3570–3575. 10.1073/pnas.1420294112

Xiong HB, Lu DD, Li ZY, Wu JH, Ning X, Lin WJ, Bai ZC, Zheng CH, Sun Y, Chi W, et al. The DELLA-ABI4-HY5 module integrates light and gibberellin signals to regulate hypocotyl elongation. Plant Commun. 2023: 4(5): 100597. 10.1016/j.xplc.2023.100597

Yan Z, Li K, Li Y, Wang W, Leng B, Yao G, Zhang F, Mu C, Liu X. The ZmbHLH32-ZmIAA9-ZmARF1 module regulates salt tolerance in maize. Int. J. Biol. Macromol. 2023: 253(Pt 4): 126978. 10.1016/j.ijbiomac.2023.126978

Yang FF, Sun YN, Du XX, Chu ZH, Zhong XH, Chen XS. Plant-specific histone deacetylases associate with ARGONAUTE4 to promote heterochromatin stabilization and plant heat tolerance. New Phytol. 2023b: 238(1): 252–269. 10.1111/nph.18729

Yang JH, Qu X, Li T, Gao YX, Du HN, Zheng LJ, Ji MC, Zhang PF, Zhang Y, Hu JX, et al. HY5-HDA9 orchestrates the transcription of *HsfA2* to modulate salt stress response in *Arabidopsis*. J. Integr. Plant Biol. 2023a: 65(1): 45–63. 10.1111/jipb.13372

Yang PC, Zhou B, Li YH. The bHLH transcription factors involved in anthocyanin biosynthesis in plants. J. Plant Physiol. 2012: 48: 747–58.

Yao SZ, Kang JR, Guo G, Yang ZR, Huang Y, Lan Y, Zhou T, Wang LY, Wei CH, Xu ZH, Li Y. The key micronutrient copper orchestrates broad-spectrum virus resistance in rice. Sci. Adv. 2022: 8(26): eabm0660. 10.1126/sciadv.abm0660

Yao XH, Fang K, Qiao K, Xiong JW, Lan J, Chen J, Tian Y, Kang XK, Lei W, Zhang DW, et al. Cooperative transcriptional regulation by ATAF1 and HY5 promotes light-induced cotyledon opening in *Arabidopsis thaliana*. Sci. Signal. 2024: 17(817): eadf7318. 10.1126/scisignal.adf7318

Yu WW, Chen QF, Liao K, Zhou DM, Yang YC, He M, Yu LJ, Guo DY, Xiao S, Xie RH, et al. The calcium-dependent protein kinase CPK16 regulates hypoxia-induced ROS production by phosphorylating the NADPH oxidase RBOHD in Arabidopsis. Plant Cell 2024: 36(9): 3451–3466. 10.1093/plcell/koae153

Yu Y, Liu HF, Xia HR, Chu ZH. Double- or triple-tiered protection: Prospects for the sustainable application of copper-based antimicrobial compounds for another fourteen decades. Int. J. Mol. Sci. 2023a: 24: 10893.

Yu Y, Xu MM, Ding XH, Chu ZH, Liu HF. Activating the *MYB51* and *MYB122* to upregulate the transcription of glucosinolates biosynthesis genes by copper ions in *Arabidopsis*. Plant Physiol. Biochem. 2021: 162: 496–505.

Yu ZP, Ma JX, Zhang MY, Li XX, Sun Y, Zhang MX, Ding ZJ. Auxin promotes hypocotyl elongation by enhancing BZR1 nuclear accumulation in *Arabidopsis*. Sci. Adv. 2023b: 9(1): eade2493. 10.1126/sciadv.ade2493

Zhang BG, Liu HF, Ding XH, Qiu JJ, Zhan M., Chu ZH. *Arabidopsis thaliana ACS8* plays a crucial role in the early biosynthesis of ethylene elicited by Cu^2+^ ions. J. Cell Sci. 2018: 131: jcs202424.

Zhang CL, Wu YJ, Liu XR, Zhang JY, Li X, Lin L, Yin RH. Pivotal roles of ELONGATED HYPOCOTYL5 in regulation of plant development and fruit metabolism in tomato. Plant Physiol. 2022a: 189(2): 527–540. 10.1093/plphys/kiac133

Zhang HY, Zhao X, Li JG, Cai H, Deng XW, Li L. MicroRNA408 is critical for the *HY5-SPL7* gene network that mediates the coordinated response to light and copper. Plant Cell 2014: 26(12): 4933–4953. 10.1105/tpc.114.127340

Zhang JH, Sun LF, Wang Y, Li BY, Li XG, Ye ZQ, Zhang J. A calcium-dependent protein kinase regulates the defense response in *Citrus sinensis*. Mol. Plant Microbe. In. 2024a:10.1094/MPMI-12-23-0208-R. Advance online publication. https://doi.org/10.1094/MPMI-12-23-0208-R

Zhang N, Hecht C, Sun XP, Fei ZJ, Martin GB. Loss of function of the bHLH transcription factor Nrd1 in tomato enhances resistance to *Pseudomonas syringae*. Plant Physiol. 2022b: 190(2): 1334–1348. 10.1093/plphys/kiac312

Zhang SX, Cao P, Xiao ZL, Zhang Q, Qiang Y, Meng H, Yang AG, An YY, Zhang MX. Rastonia solanacearum type Ⅲ effectors target host 14-3-3 proteins to suppress plant immunity. Biochem. Bioph. Res. Co. 2024b: 690: 149256. 10.1016/j.bbrc.2023.149256

Zhao L, Xie L, Zhang Q, Ouyang WZ, Deng L, Guan PP, Ma M, Li Y, Zhang Y, Xiao Q, et al. Integrative analysis of reference epigenomes in 20 rice varieties. Nat. Commun. 2020: 11(1): 2658. 10.1038/s41467-020-16457-5

Zhao XB, Wang Q, Yan CX, Sun QX, Wang J, Li CJ, Yuan CL, Mou YF, Shan SH. The bHLH transcription factor AhbHLH121 improves salt tolerance in peanut. Int. J. Biol. Macromol. 2024: 256(Pt 2): 128492. 10.1016/j.ijbiomac.2023.128492

Zhao Y, Shi H, Pan Y, Lyu M, Yang ZX, Kou XX, Deng XW, Zhong SW. Sensory circuitry controls cytosolic calcium-mediated phytochrome B phototransduction. Cell 2023: 186(6): 1230–1243.e14. 10.1016/j.cell.2023.02.011

